# Bio-engineering a common probiotic to exploit colonic inflammation promotes reliable efficacy in translational models of colitis

**DOI:** 10.1101/2024.10.08.617317

**Authors:** Andrea Verdugo-Meza, Sandeep K. Gill, Artem Godovannyi, Malavika K. Adur, Jacqueline A. Barnett, Mehrbod Estaki, Jiayu Ye, Natasha Haskey, Hannah Mehain, Jessica K. Josephson, Ray Ishida, Chanel Ghesquiere, Laura M. Sly, Deanna L. Gibson

## Abstract

The intricate balance between the gut microbiome and host health inspires innovations in drug development. Commensal bacteria provide a multi-targeted approach ideal for treating complex medical conditions, like inflammatory bowel disease (IBD). These bacteria are self-replicating factories with broad targets that promote balanced intestinal inflammation, mucosal barrier function, and eubiosis. Yet, the lack of superiority to gold-standard treatments and their clinical inconsistency makes most probiotics unreliable for disease treatments. Intestinal inflammation, a driving factor in many diseases, often overwhelms commensal bacteria, which lack the stress-resistance mechanisms necessary to withstand host immune defenses. To address this, we introduced a persistence platform BioPersist™ into *E. coli* Nissle 1917. We hypothesized that a bio-engineered probiotic, or genetically engineered microbial medicine (GEMM™), designed to persist during inflammation would enhance probiotic bioavailability during colitis, leading to sustained therapeutic outcomes. We evaluated BioPersist in multiple translational colitis models such as in mice and pigs. BioPersist delayed the onset and reduced the severity of both chronic and acute colitis, proving more effective than 5-aminosalicylate. BioPersist thrived during inflammation promoting tolerogenic immune responses that limited infiltrating leukocyte activity and decreased TNF-α from resident myeloid cells in the mesentery. The persistence feature of BioPersist allowed the probiotic to overcome the damaging inflammatory response, eliciting mucosal healing evident by the increase in microbially-derived butyric acid. Based on these preclinical results, BioPersist may be a novel therapeutic option for both human and veterinary applications that sustains efficacy during colitis.

**One Sentence Summary:** Adding a persistence feature to a probiotic enhances its efficacy for colitis treatment, enhancing future human and veterinary therapeutic applications.

## INTRODUCTION

Many modern diseases, such as inflammatory bowel disease (IBD), metabolic syndrome and neuropsychiatric disorders, have been linked to chronic inflammation in the gastrointestinal tract. Unlike conventional drugs, which target a single pathway or molecule, health-promoting bacteria provide a multi-targeted approach desirable for biologically complex conditions. Beneficial bacteria can modulate inflammation, strengthen the intestinal barrier, and reestablish host-microbiome symbiosis *(1, 2)*. While the use of probiotics has been promoted to induce health benefits in humans and domestic animals for decades, objective clinical outcomes are inconsistent. In the IBD population, several meta-analyses conclude that probiotics are not clinically effective *(3, 4)*, while others show only patients in remission receive benefits *(5)*. This clinical inconsistency may be related to the fact that probiotics only have transient beneficial responses, and that they may only be effective in patients in remission. A key reason for this is the inconsistent engraftment of probiotics in the gut, which limits their ability to provide sustained therapeutic benefits. Despite consumption, probiotics often fail to establish stable colonization *(6)*, and their persistence after cessation is highly variable *(7)* and generally low *(1)*. Even when probiotics possess factors that aid adhesion, their effects are short-lived as many do not remain in the gut long-term. For instance, commonly used probiotics including the bacterial consortium VSL#3 *(8, 9)*, *Lactobacillus sp*.*(10)*, and Mutaflor® (EcN) *(11, 12)* have been shown to lack sustained persistence in the gut.

Another factor impacting probiotic persistence and robustness is the significant oxidative stress present during inflammation *(13-16)*. Elevated levels of reactive oxygen species in the inflamed mucosa suppress the growth of symbiotic bacteria in the bowel *(17, 18)*. It is plausible that the poor clinical efficacy of many probiotics is due to insufficient bioavailability in the inflamed IBD intestine. Chronic inflammation modifies the intestinal ecosystem, creating a niche that favors the colonization of pathobionts over commensal microbes *(19)*. It is unclear if commensals are depleted in the IBD intestine due to inflammation or if a lack of commensal microbes predisposes to the development of IBD. While no universal gut microbiota profile is consistently observed at diagnosis, evidence supports a general deficiency of beneficial commensal microbes and an overgrowth of harmful pathobionts. Unlike beneficial microbes, many pathogens successfully exploit host defenses as they have evolved stress survival systems that give them the fitness to thrive in unfavorable conditions *(19)*. *Salmonella enterica* serovar Typhimurium strain succeeds during inflammation due to its ability to utilize tetrathionate (TTR), a metabolite produced in the inflamed intestine, as a terminal electron acceptor *(17)*. Thus, the tetrathionate operon *(ttr* operon) may act as a genetic feature that supports bacterial expansion in the IBD intestine.

An emerging area of innovation in microbiome targeted therapies is the use of live biotherapeutics, which differ from over-the-counter probiotics given their intended use as evidence-based treatments studied for specific ailments. These include bacteria, genetically engineered to enhance their beneficial health effects, or genetically engineered microbial medicines (GEMM™). To overcome the natural limitations of probiotics’ low persistence in the inflamed intestine, we bioengineered a novel GEMM, designed to enhance the persistence of *Escherichia coli* Nissle 1917 (EcN) in the inflamed bowel *(20)*. Bio-engineered persistent EcN (referred to here as BioPersist™) has a chromosomal insert of the *ttr* operon. We postulated that enhancing EcN with persistence features would improve its bioavailability and provide sustained efficacy during colitis *(20)*. In support of this, an exploratory study of the colon metabolome revealed that BioPersist promotes the secretion of unique gut-derived metabolites during DSS-colitis, unlike the parental strain EcN *(21)*. In this study, we show BioPersist blooms during colitis and, unlike EcN, can persist long-term in a model of chronic colonic inflammation (Muc2^- /-^ mice). BioPersist delays and protects against the development of severe colitis by reducing pro-inflammatory immune responses involving key inflammatory cell populations while promoting the expansion of T-regulatory cells and the induction of a key commensal metabolite, butyric acid, with anti-inflammatory effects. The protection elicited by the GEMM was translatable from colitis rodent models to a porcine model. In summary, pre-clinical studies provide evidence that BioPersist is a reliable therapeutic with sustained efficacy during colitis. Future considerations to create novel GEMMs using the BioPersist persistence platform could enhance microbiome-based therapeutics’ potential for human and animal applications.

## RESULTS

### BioPersist blooms during inflammation in the large bowel

Bio-engineered persistent EcN (referred to here as BioPersist; Fig. 1A) was genetically modified to carry the *ttr* operon chromosomally, and we hypothesized this would increase its persistence and, therefore, clinical efficacy during colitis *(20)*. To support this concept, we evaluated EcN and BioPersist longitudinal detection in healthy (C57BL/6 mice) and chronically inflamed colitic mice (Muc2^-/-^ mice). As observed in Fig. 1B, BioPersist was detected for an extended period (up to 22 weeks post-treatment; Fig S1) compared to Muc2^-/-^ mice receiving the parental strain EcN, which was mostly undetectable. Unlike the Muc2^-/-^ colitis-prone mice, healthy C57BL/6 WT mice harbored little to no BioPersist and displayed no observable histological, physiologic, or immunologic changes (Fig. S2). We confirmed that BioPersist bloomed in colitic Muc2^-/-^ mice by correlating the detection levels of *ttr* copies with the clinical scores (Fig. 1C). Notably, BioPersist blooms can be observed following increases in clinical scores and recedes once clinical scores decrease (○ = grey follows *ttr* copies and □ = blue line follows clinical scores). This data supports our hypothesis that the *ttr* operon helps EcN bloom in the inflamed colon and loses its ability to dominate the healthy gut. Additionally, 16S rRNA gut microbiome data shows that BioPersist does not over-colonize the gut following the intervention, with less than 5% overall present in the ecosystem (Fig. S3).

**Figure 1.**
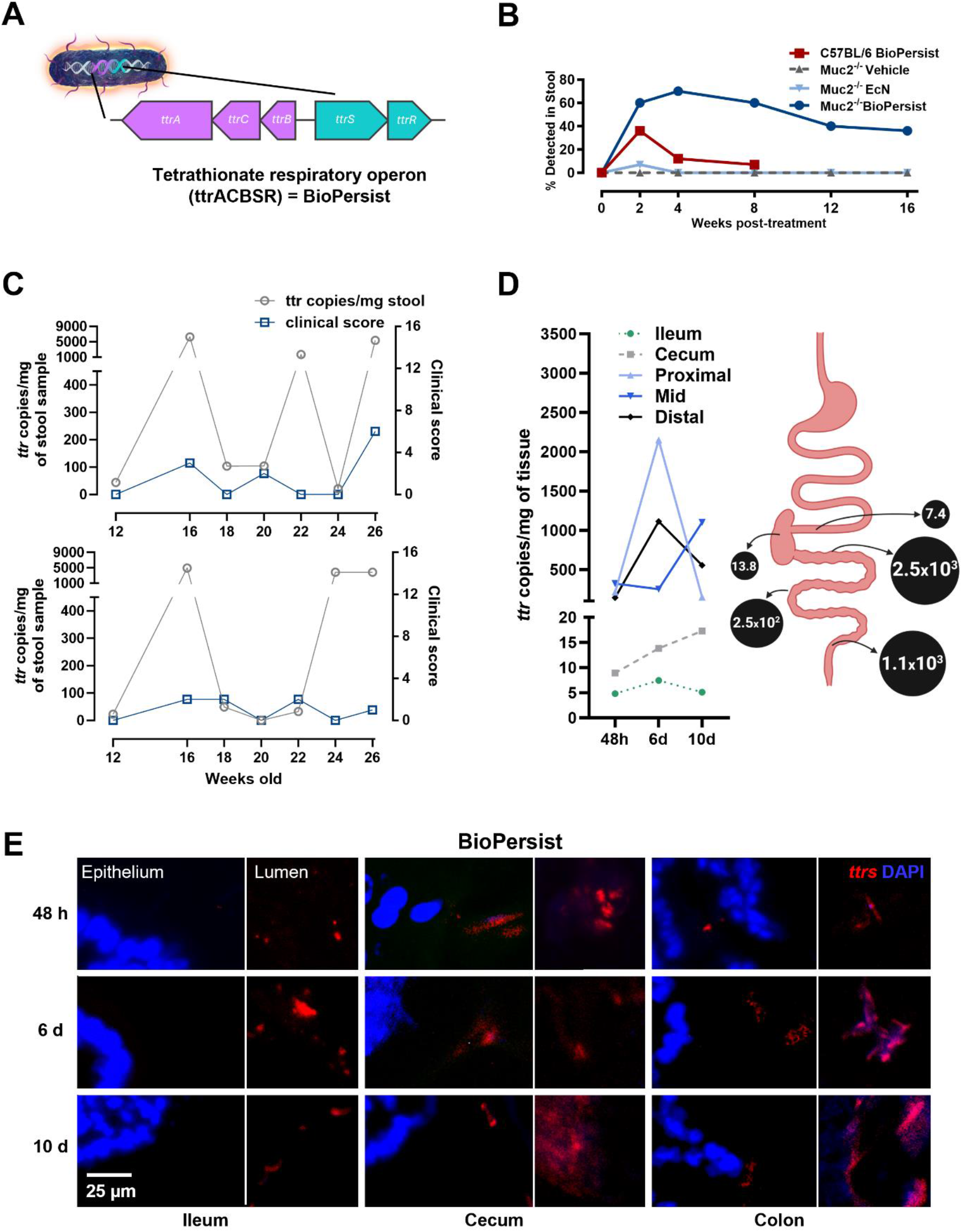
BioPersist thrives under colitis conditions. (**A**), genetic modification of BioPersist; (**B**), persistence of BioPersist in stool samples of colitic (Muc2^-/-^ mice) and healthy mice (C57BL/6 mice); (**C**), correlation of BioPersist load *(ttr* copies/mg of stool) against clinical scores of colitis in mice; (**D**), detection of BioPersist along the gastrointestinal tract; (**E**), visualization of BioPersist via FISH staining in the gastrointestinal tract. The results were obtained from Muc2^-/-^ mice except in (B) where C57BL/6 were also included. N = 3-11.

To understand where BioPersist preferentially occupies a niche in the IBD gut, we examined its presence from the ileum to the distal colon. After a single orally administered dose to Muc2^-/-^ mice, we found BioPersist occupied all regions of the large bowel and was observed close to the epithelial surface (Fig. 1D-E). In contrast, the ileum harbored little BioPersist notably absent near the mucosal surface. To clarify if BioPersist could still influence disease activity in the ileum, we examined SHIP^-/-^ mice (mice deficient in Src homology 2 domain-containing inositol 5’-phosphatase), a model of Crohn’s disease-like inflammation in the ileum *(22)*. Consistent with the finding that BioPersist was undetectable in the ileum, BioPersist did not affect ileitis (Fig. S4). These findings suggest that the probiotic’s benefits are associated with its presence in the region of colonization. Overall, these data reveal BioPersist has a preferential colonization for the large bowel during colitis.

### BioPersist is more effective than front-line therapy 5-ASA in acute colitis

The global disease burden of IBD, including ulcerative colitis (UC) and Crohn’s disease, is significant *(23)*, with prevalence continuing to increase *(24)*. The front-line treatment for UC has been the administration of aminosalicylates (5-ASA) *(25)*; but as with other IBD drug treatments, sustained remission and complete mucosal healing has remained elusive. First-line therapies like 5-ASA provide temporary benefits in mild-moderate IBD but poor patient acceptability, adherence, and frequent dosing *(26)* limit its clinical usefulness. In addition, 30% of UC patients are refractory to 5-ASA *(27)*, and within the first year, a further 40% of patients receiving the biologic infliximab will loss responsiveness *(28-30)*. To determine how BioPersist compares to a front-line therapy, we compared its efficacy with 5-ASA in the dextran sodium sulfate (DSS) induced acute colitis model, which models an acute flare in UC *(31)*. We exposed mice to 3.5% DSS in their drinking water for seven consecutive days post-intervention (Fig. 2). Reported effectiveness of 5-ASA has been variable, with some studies showing a reduction of colitis, while others have reported it has little effect *(32)*. We included several controls to compare the efficacy of BioPersist against no treatment (vehicle group) and treatment with EcN (the unmodified probiotic strain). Mice were monitored daily for disease activity which included scores for diarrhea, dehydration, lethargy and weight loss. Only mice treated with BioPersist presented significantly lower clinical scores (Fig. 2A), while both EcN and BioPersist-treated mice showed reduced weight loss (Fig. 2B). The drug 5-ASA had little effect in protecting mice from DSS colitis, a finding already reported *(32)*. When we evaluated the macroscopic changes amongst the groups, we found that only mice treated with BioPersist had formed stool pellets in the distal colon and had a longer colon length (Fig. S5). BioPersist-treated mice had the lowest pathology (Fig. 2C) calculated from histopathological scores (Fig. 2D) characterized by less epithelial erosion, more crypts with goblets cells and lower immune cell infiltration. We further examined leukocyte infiltration by identifying infiltrating macrophages and neutrophils, known drivers of UC and evident in many colitis’ models. Again, EcN and BioPersist treatment appeared to reduce macrophage and neutrophil infiltration, but only BioPersist presented significant differences (Fig. 2E). We then investigated how BioPersist modulates short-chain fatty acid production since this is a key beneficial response from the gut microbiome and found an increase in acetic and butyric acid, with a reduction in propionic acid (Fig 2F). Other factors that we explored and found specifically modulated by the persistence feature of BioPersist were decreased levels of fecal calprotectin (Fig. 2G) and a reduction in the relative expression of *lcn2* (lipocalin 2) (Fig. 2H), both clinical markers of neutrophil activity. On the other hand, BioPersist induced greater expression of *Il22* (IL-22) in this model (Fig. 2I), an important cytokine implicated in mucosal healing *(33)*. Modest upregulation of other genes involved in mucosal healing, including antimicrobial peptides and mucin-2 were also observed with BioPersist treatment (Fig. S5). Overall, BioPersist was consistently more efficacious during acute colitis, unlike its unmodified form EcN and front-line therapy 5-ASA.

**Figure 2.**
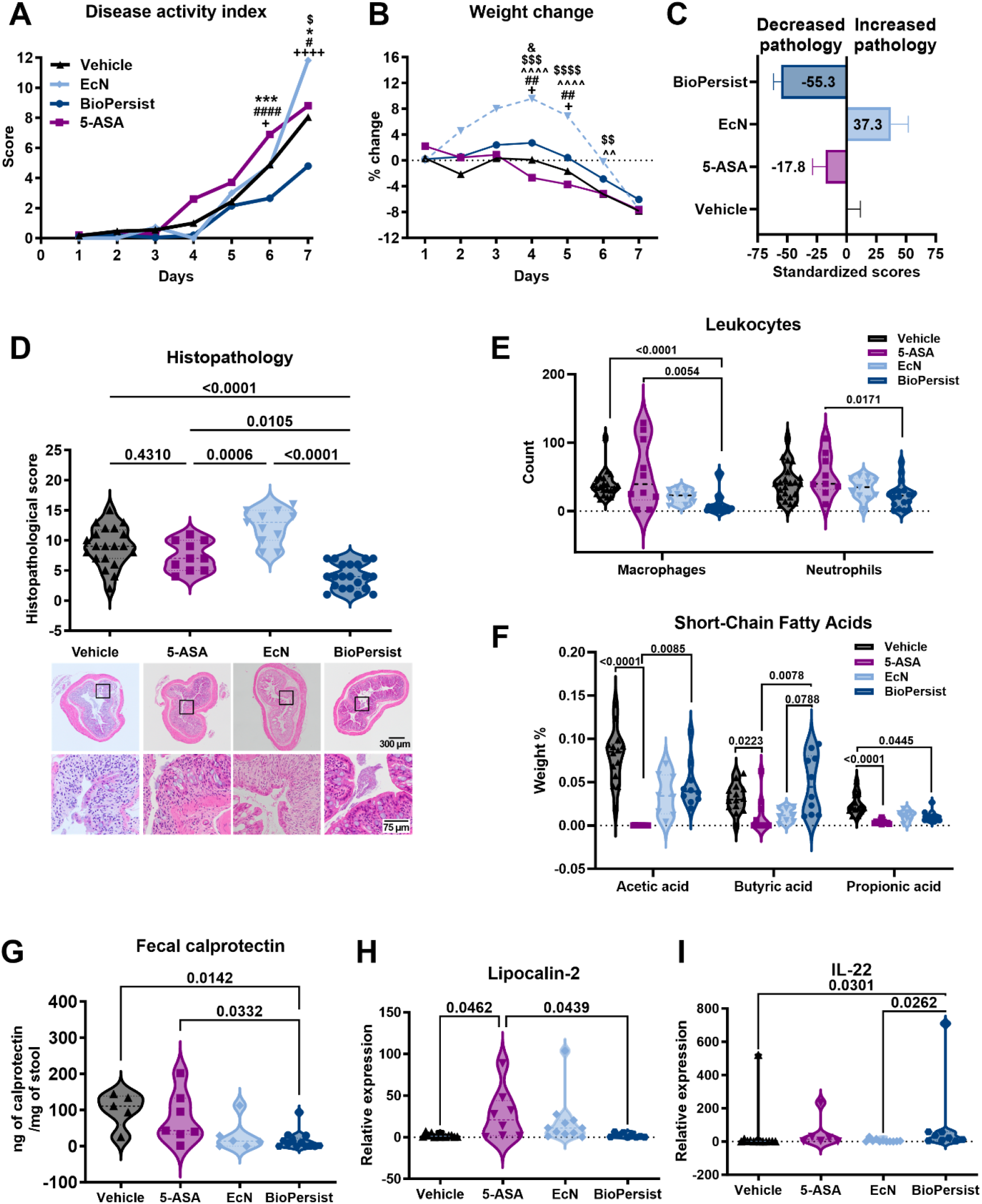
BioPersist protects more effectively than front-line 5-ASA therapy in the DSS model of colitis. (**A**), overall disease activity index, (**B**), daily weight changes; (**C**), standardized histopathological scores to vehicle group’s scores showing decreased (left) or increased (right) pathology. (**D**), histopathological scores and representative H&E-stained section of the distal colon; (**E**), leukocyte infiltration counts in distal colon sections stained with fluorescently labeled antibodies against F4/80 (macrophages) or MPO (neutrophils); (**F**), short-chain fatty acid content in cecum; (**G**), Fecal calprotectin quantification; (**H**) and (**I**), relative expression of lipocalin 2 *(Lcn2)* and IL-22 *(Il22)* in the distal colon. N = 10-20 mice per group. Data was analyzed with One-way ANOVA (A, B, D) or Kruskal-Wallis (E-I). Adjusted *P*-values are shown in graphs.

### BioPersist delays and protects against chronic and spontaneous colitis via modulation of tolerogenic immune networks

To further understand the potential therapeutic effect of BioPersist during chronic inflammation in the colon, the site of its expansion, we observed Muc2^-/-^ mice long term. Young mice with subclinical colitis received BioPersist and were compared to mice gavaged with vehicle (media) or the parental probiotic EcN. After the initial dosing of bacteria, the mice were monitored weekly until they were at least four months old, when they frequently develop rectal prolapses, the most severe sign of disease in the Muc2^-/-^ model *(34)*. Clinical scores were lower for mice treated with BioPersist (1.0±1.19) compared to mice treated with vehicle or EcN (2.86±2.19) (Fig. 3A). Contrary to the 11-30% rectal prolapses recorded in the vehicle or EcN group, BioPersist-treated mice did not develop any rectal prolapse by this timepoint (Fig. 3B), although by 8 months of age, these mice eventually did develop rectal prolapses, albeit at a much-reduced frequency compared to the controls (Fig. S6; 40% BioPersist vs 60-80% controls). Consistent with these findings, colons were longer in mice treated with BioPersist (Fig. 3C), a sign of reduced colitis. The histopathology assessment confirmed our clinical observations. Among the three groups, mice treated with BioPersist had the lowest histopathological scores, characterized by decreased epithelial damage and preserved crypt architecture (Fig. 3D).

**Figure 3.**
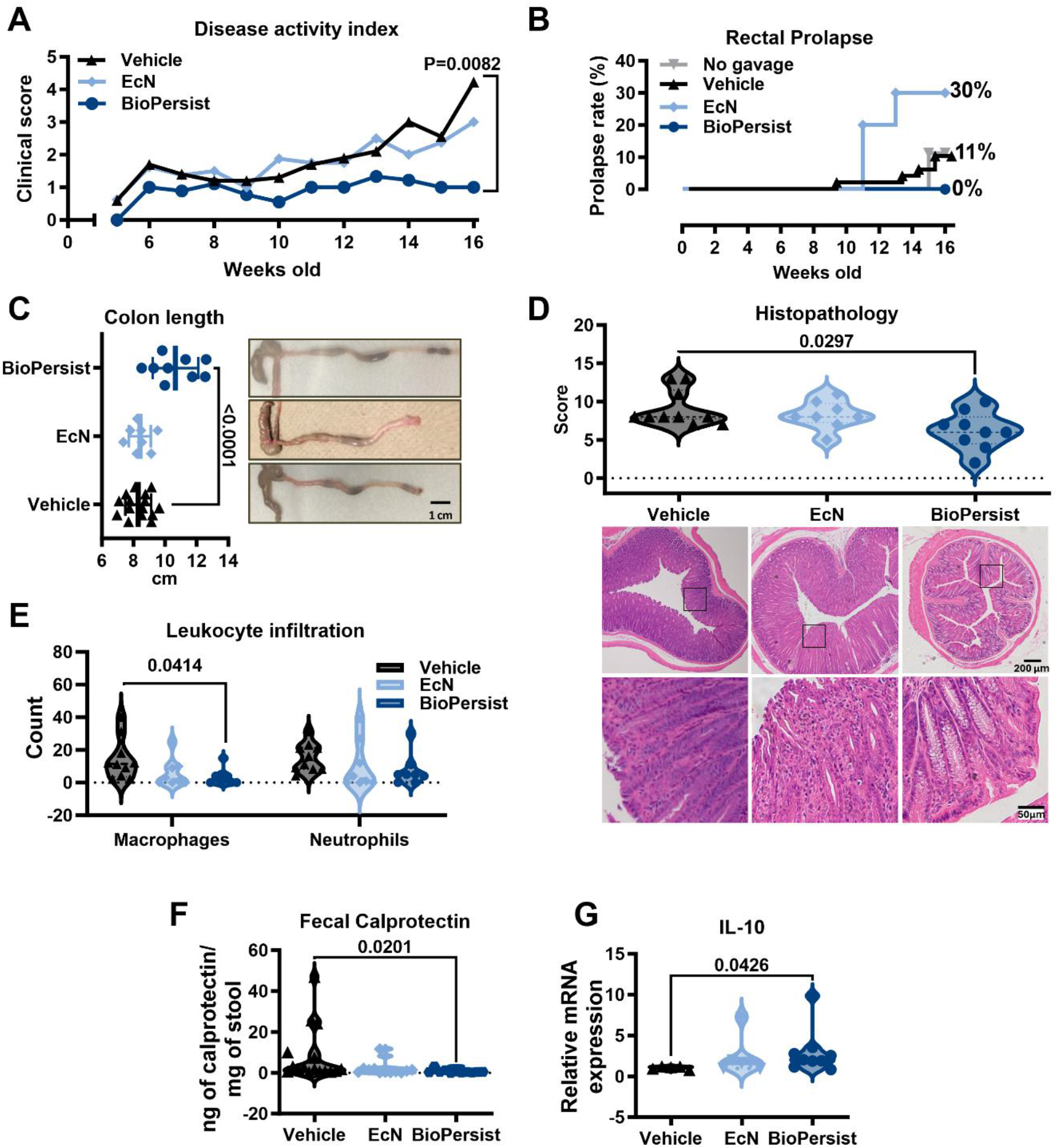
BioPersist decreased and delayed the colitic phenotype of Muc2^-/-^ mice. (**A**), disease activity index shows the rise in clinical scores starting at 6 weeks of age; (**B**), Rectal prolapse rate; (**C**), macroscopic changes and colon length. (**D**), Histopathological scores and representative H&E sections of the distal colon; (**E**), leukocyte counts in distal colon sections stained with fluorescently labeled antibodies against F4/80 (macrophages) or MPO (neutrophils); (**F**), Fecal calprotectin quantification when mice were 3 months old; (**G**), relative mRNA expression of IL-10 *(Il10)* distal colon of 3-month-old mice. N = 5-23 mice per group. Data was analyzed with One-way ANOVA (A, C, and D), Kruskal-Wallis (E-G), or Log-rank (Mantel-Cox) test (B). Adjusted *P*-values are shown in the graphs.

As Muc2^-/-^ mice are constantly challenged by luminal microbial triggers due to their absence of the protective mucin-2 layers, their immune response is always active, driving the damage observed in their colons. BioPersist-treatment resulted in fewer infiltrating macrophages and neutrophils (Fig. 3E) associated with reduced fecal calprotectin at 12 weeks of age (Fig. 3F). Despite BioPersist delaying severe onset of colitis in this model, between 14-22 weeks post-intervention, signs of progressive colitis were evident, albeit to a lesser extent (Fig. S6). We also observed that BioPersist increased relative expression of *Il10* (Fig. 3G), a gene encoding the anti-inflammatory cytokine IL-10. Therefore, we decided to explore the immunomodulatory effect of BioPersist in the gut microenvironment. We looked at different markers of the immune response, from antimicrobial peptides to cytokines, and we found a general downregulation of most of these genes (Fig. 4A). It is well reported in the Muc2^-/-^ model that increased expression of the antimicrobial peptide RELM-β correlates with rectal prolapse and severe disease *(34)*. We found that the relative expression of RELM-β was significantly lower in mice treated with BioPersist. A similar trend was found for other antimicrobial peptides (RegIII-γ, RegIII-β, mMBD2, mMBD3, mMBD4 and mDef6), pro-inflammatory cytokines (IL-1β, IFN-γ, IL-17, IL-6), barrier function proteins (ZO-1 and ZO-2) and innate immune recognition (TLR-4). We immunophenotyped the colonic lamina propria cells by flow cytometry; ultimately, these cells orchestrate the immune response (Fig 4B-I and S7-8). Mice in the BioPersist group had lower % of eosinophils (Fig. 4B) and neutrophils (Fig. 4C) but higher % of T-regulatory cells (Fig. 4D). For cell secreting cytokines, we found lower levels of macrophages (Fig. 4E) secreting TNF-α and lower levels of leukocytes secreting IL-22 (Fig. 4F) 16 weeks post-intervention with BioPersist. While the % of dendritic cells were similar (Fig. 4G), there was a decrease in the pro-inflammatory phenotype of other resident myeloid cells in the lamina propria producing TNF-α (Fig. 4H-I), suggesting that BioPersist promotes a more tolerogenic effect during chronic colitis, where resident myeloid cells and T-regulatory cells may suppress activation and infiltration of macrophages, neutrophils and eosinophils.

**Figure 4.**
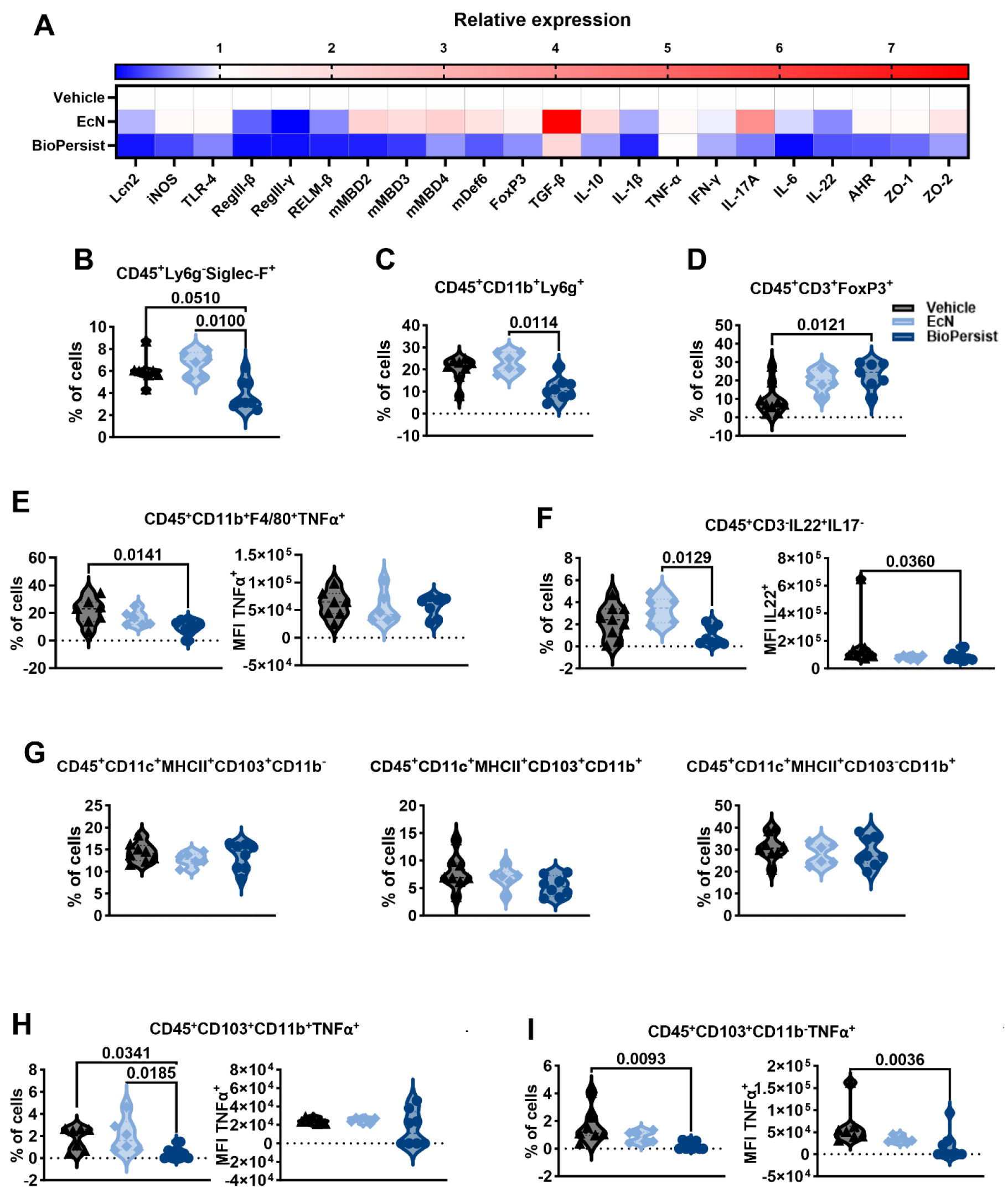
BioPersist promotes a tolerogenic immune response in Muc2^-/-^ mice. (**A**), relative mRNA expression of different cell markers, antimicrobial peptides, cytokines, transcription factors, and barrier proteins in the distal colon; (**B** to **I**), immunophenotyping of colonic lamina propria cells via flow cytometry; (B), eosinophils; (C), neutrophils; (D), T-regulatory cells; (E), macrophages secreting TNF-α; (F), leukocytes producing IL-22; (G), dendritic cells; (H-I), resident leukocytes secreting TNF-α. N = 6-8. Data was analyzed via Kruskal-Wallis and adjusted *P-*values are expressed in each graph. MFI = Mean Fluorescence Intensity.

### The protective effect of BioPersist is translated in larger mammals

Pigs and humans share a more similar gut anatomy and physiology and, pigs are also highly susceptible to gastrointestinal diseases *(35)*. To advance in translating BioPersist into a clinical treatment, we evaluated the therapeutic effect of BioPersist in pigs with chemically induced colitis. Pigs were exposed to a half dose of DSS and then orally received a single dose of BioPersist or a vehicle control followed by daily oral gavage of DSS for 7 days. BioPersist-treated pigs displayed increased weight gain and had a predicted probability of increased average daily gain (ADG) when compared to pigs in the vehicle control cohort (Fig. 5A). Additionally, BioPersist positively influenced survival rates (Fig. 5B) by preventing mortality, unlike the vehicle control group where ∼20% mortality was observed. Rectal temperature was recorded as a sign of disease onset, which showed that BioPersist treatment protected from high fever (Fig. 5C). Necropsy findings indicated minimal macroscopic damage in the BioPersist cohort, contrasting sharply with the enlarged, bloody colons observed in the vehicle control group (Fig. 5D). Consistent with these observations, histopathological images revealed fully formed crypts and visible goblet cells with little epithelial erosion unlike the vehicle control colitis group (Fig 5E) corresponding to reduced scores for damage (Fig 5F). As DSS-induced colitis leads to bacterial translocation due to a severely compromised intestinal barrier, we compared the bacterial load in blood, spleen and mesenteric lymph nodes (MLN). Critically, BioPersist was not found to translocate into blood, MLN or the spleen when samples were plated on selective media containing antibiotics (Fig 5G-I). Instead, BioPersist treatment reduced the generalized bacterial translocation to the spleen and MLN when compared to pigs in the vehicle control group (Fig. 5E). Overall, BioPersist protected pigs with an intestine that structurally and physiologically is more representative of the human intestine, from severe chemically induced colitis supporting a possible beneficial role in humans.

**Figure 5.**
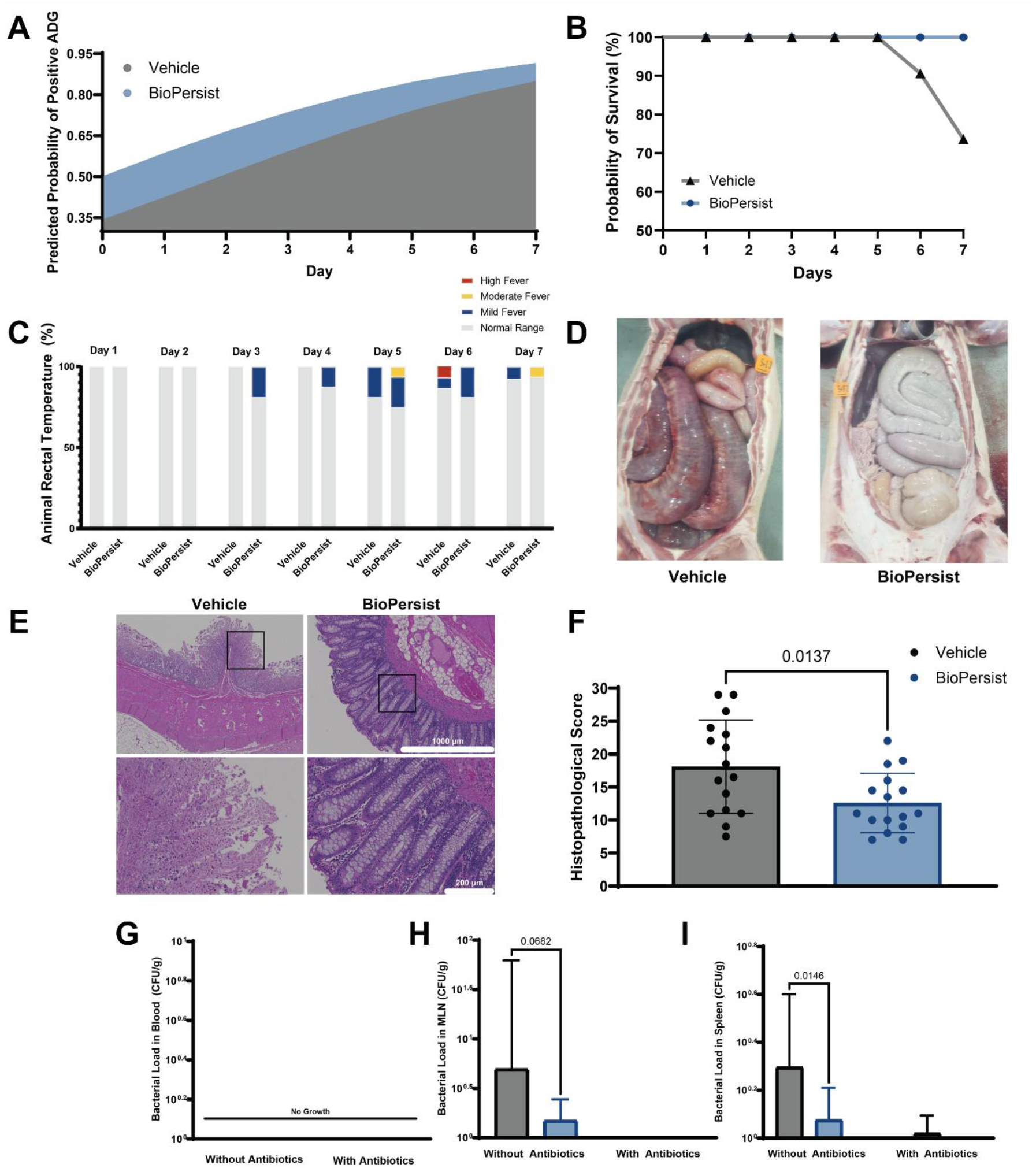
Pigs are also protected from severe DSS-induced colitis when introduced with BioPersist. (**A**), the probability of positive ADG after exposure to DSS. (**B**), probability of survival. (**C**), rectal temperature changes through colitis induction. (**D**), representative images of the macroscopic changes in the colon. (**E**), representative colon sections. (**F**), histopathological scores. (**G** to**I**), bacterial load in blood, MLN and spleen, respectively. N = 16 pigs per group. Predicted probability was determined using a binary generalized linear model. Adjusted *P-*values are expressed in each graph. ADG = average daily gain; MLN = mesenteric lymph nodes.

## DISCUSSION

In this study, we demonstrate that a GEMM developed using the BioPersist platform can overcome the limitations of its natural parent probiotic, EcN, through the introduction of a persistence feature that enables it to thrive during colitis. In head-to-head comparisons, BioPersist outperformed both the unmodified EcN strain and the current gold-standard treatment for mild-moderate UC, 5-ASA, showing significant protective effects across multiple rodent and porcine models of colitis. Sustained efficacy was concomitant with a reduction in clinical biomarkers like fecal calprotectin, the induction of key mucosal anti-inflammatory molecules including butyric acid, and the shift from a pro-inflammatory immune milieu to a tolerogenic gut microenvironment, with the modulation of key immune cell populations including the induction of T-regulatory cells (summary Fig. 6). Overall, we provide pre-clinical evidence that adding the persistence feature to EcN (BioPersist) results in a reliable and effective therapeutic for colitis with clinical applications. Modifying commensal bacteria to specifically thrive in the IBD gut could improve their ability to elicit their beneficial effects including mucosal healing.

**Fig. 6.**
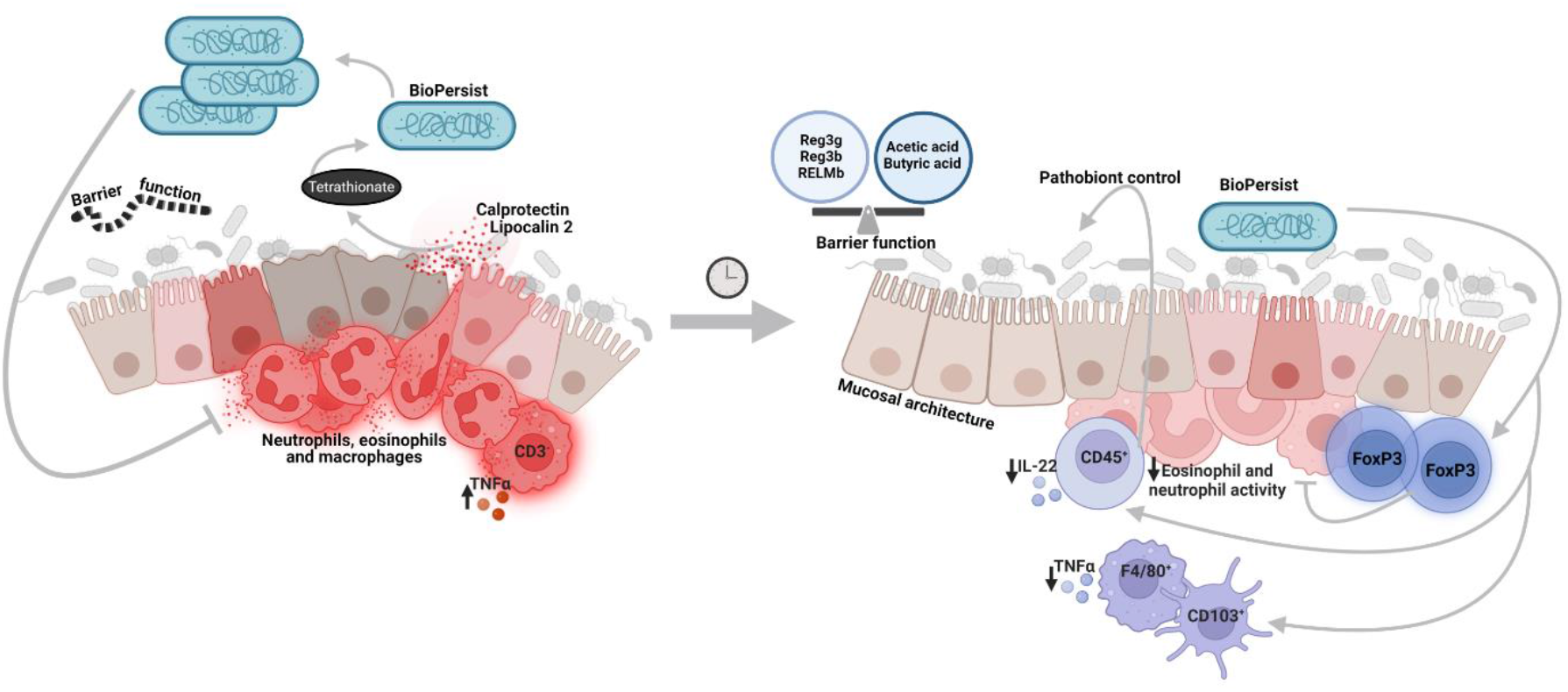
Working model for BioPersist in the colitic colon. The inflamed colon favors BioPersist to thrive and impact the microenvironment by balancing barrier and barrier-related molecules, preventing pathobiont growth, and by promoting a tolerogenic environment characterized by T-regs and leukocyte with low TNF-α expression that is associated to decreased presence of eosinophils, macrophages and neutrophils in the lamina propria of the colon. Figure created in Biorender.

Probiotics can be considered natural factories creating numerous beneficial compounds, yet interventions have not translated into clinically relevant results for complex diseases like IBD. Accounting for the limited survival of probiotics in the GI tract, we hypothesized that a hostile microenvironment present during colitis impedes probiotics to persist and have a therapeutic impact. As we show here, the probiotic EcN is eradicated in the colitic microenvironment during chronic colitis in Muc2^-/-^ mice, similar to what has been observed in humans *(11)*. However, when the probiotic was genetically engineered to carry a persistence feature, BioPersist was found to remain present and even bloomed during bouts of flares in the progressive chronic Muc2^-/-^ colitis model. While BioPersist was effective in reducing colitis, it did not protect against ileitis, consistent with a preference to occupy the large but not the small bowel. This is a key observation since it demonstrates that the absence of the probiotic in the ileum corresponded to a lack of disease protection. This contrasts with the concept of “paraprobiotics” claiming non-viable probiotics retain their beneficial effect *(36)*. Based on our findings, we hypothesize that viability and presence in the site of disease are correlated. Indeed, BioPersist persisted in the large bowel correlating with protection related to improvement of the intestinal barrier evident with the lack of bacterial translocation in the porcine tissues. BioPersist also increased RegIII-γ, which can help restructure the intestinal microbiome for clearance of harmful pathogens reducing chronic inflammation. BioPersist was able to re-program immune cell development resulting in increases in T-regulatory cells that protected mice against developing severe colitis. Pro-inflammatory immune networks involving granulocytes such as eosinophils, macrophages and neutrophils were downregulated resulting in lower fecal calprotectin and lipocalin2. Although it has been reported that EcN decreases CD45^+^ cell infiltration in the DSS model of colitis *(37)*, our results show only BioPersist and not EcN, can provide this therapeutic outcome in a chronic model, which is more translatable to human UC. This is despite our minimal intervention with doses administered only during the early stages of disease in Muc2^-/-^ mice, compared to the reported literature for DSS models where mice must receive daily doses of EcN *(37)*.

Typical therapy for IBD includes immunosuppression which leads to risk of infectious disease and cancer *(38, 39)*. EcN has long been recognized to protect from infectious pathogens which implies this bacterial therapy does not shut down critical defensive immunity but instead regulates the gut microenvironment to be more tolerogenic during chronic colitis. Indeed, EcN modulates tight junction proteins, antimicrobial peptides, cytokines and other factors involved in defensive immunity *(40)*. For example, Sturn et al. showed *in vitro* that T cells derived from the intestine limit their expansion when exposed to EcN supernatant, while peripheral T cells are unaffected *(41)*. Regarding EcN in models of colitis, mice exposed to DSS and receiving oral gavage with EcN show milder colitis dependent on Toll-like receptor signaling *(42)*. However, growth and survival of EcN and other probiotics is compromised in an inflammatory environment, shortening the organisms’ beneficial effects. In contrast, BioPersist has the “fitness tools” to thrive under inflammatory conditions, allowing sustained therapeutic benefits. Here we show what others in the literature have demonstrated, mice receiving EcN had a delayed onset of DSS-colitis, decreased leukocyte infiltration, and reduced inflammation. However, like findings from clinical trials, the overall endpoint results for the EcN group were not better than those found in the control group. Our results in Muc2^-/-^ mice support the lost therapeutic effect, where only the treatment with BioPersist decreased the severity of colitis 8 weeks post-intervention and beyond. Although EcN-treated mice shared some outcomes with BioPersist-treated mice, there were important aspects where EcN-treated mice did as poorly as vehicle-treated mice. For example, while 4 weeks post-intervention, Muc2^-/-^ mice treated with EcN showed reduced fecal calprotectin and IL-10, by 8 weeks post-intervention, differences between EcN- and BioPersist-treated mice became striking, shown by the large differences in rectal prolapse rates between the two cohorts, revealing the lack of EcN persistence results in transient and inconsistent effects.

The opportunity for clinical translation of BioPersist in mild to moderate UC has the potential to initiate a tolerogenic immune balance. However, the heterogenicity of intestinal microbes and immune responses amongst individuals with complex diseases like IBD needs to be considered. While EcN has been the protagonist of biotherapeutics for cancer and metabolic therapies because of its historically reported safety record, there may be limitations with exclusively relying on EcN for next-generation biotherapeutics. With the increased high throughput of personalized omics, we anticipate incorporation of the persistence feature to create other GEMMs tailoring the therapy to suit personalized medicine. For example, new studies are shedding light on individualized microbiome changes that, along with known susceptibility to IBD *(43)*, could prompt a re-introduction of the microbes that are being lost but now equipped with a fitness advantage to persist, restoring host-microbe symbiosis. As seen here with EcN, the persistence of probiotics enhances their efficacy and longevity of their health-promoting effects. Herein lies a great potential of applying this approach to other probiotics to treat the myriad of human and animal conditions associated with intestinal inflammation.

## MATERIALS AND METHODS

### Study design

The study aimed to determine the efficacy of bioengineered EcN (BioPersist) in protecting small and big mammals from severe colitis and how this correlated with persistence in the intestine. BioPersist was compared to no treatment (vehicle group) and the unmodified probiotic (EcN) in acute DSS-induced colitis in rodents and pigs and chronic colitis in mice using a Muc2-/-model. Animals were allocated randomly to control or experimental groups repeated independently by two separate researchers. All the protocols involving the care and handling of the mice were approved and overseen by the University of British Columbia’s Animal Care Committee per guidelines by the Canadian Council on Animal Care. Mice were excluded if they had fighting lesions, extensive barbering, malocclusions, signs of illness or death unrelated to the study. The rodent studies were semi-blinded. All the protocols involving the care and handling of pigs were approved by the institutional animal care and use committee and overseen by ClinVet in South Dakota. Pigs were excluded if they showed signs of lameness, illness or death unrelated to the study. The pig trial was carried out in a double blinded manner. The sample size for mice and pigs was based on power analysis with alpha=0.05, power =0.9, and effect size from previous studies and experience. Sample processing and data acquisition were performed blindly.

### Bacterial strains and culture conditions

*Escherichia coli* Nissle 1917 (EcN) was isolated from the commercial formulation Mutaflor. The genetically engineered microbial medicine (GEMM) BioPersist was generated as described previously *(20)*. Briefly, BioPersist or a version of BioPersist (1177-00-P01-US-PRV) was engineered by cloning in a chromosomal insert of the *Salmonella enterica ttr* operon into EcN and confirmed by growth and sequencing. Bacteria were cultured in Luria-Bertani broth (per 1 L of medium: 10 g of tryptone, 5 g of yeast extract, and 10 g of NaCl, pH = 7.5) under agitation at 180 rpm at 37°C.

### Experimental animal models, housing and diets

Rodent experimental protocols were approved by the University of British Columbia’s Animal Care Committee, per guidelines drafted by the Canadian Council on the Use of Laboratory Animals under protocols A15-0201, A19-0286 and, A23-0033. C57BL/6 mice (Charles River; Wilmington, Massachusetts, USA); C57BL/6 and Muc2^-/-^ mice *(44)* were maintained under specific pathogen-free conditions, with controlled temperature at 22±2°C, 12h light/dark cycles, and providing water and food *ad libitum*. C57BL/6 mice, 4-5 weeks old, were acclimatized for a week prior to breeding or experimentation. All Muc2^-/-^ mice were bred in-house (Animal Care Center at the University of British Columbia Okanagan, Kelowna, Canada) fed normal chow or breeding chow (PicoLab Mouse Diet 20: 5053), *ad libitum* and autoclaved water. Both male and female mice were included in each experiment, and we did not observe any statistical differences between sexes thus, the groups were pooled for analysis.

The pig trial was carried out by a contract research organization, ClinGlobal ClinVet in South Dakota https://clinvet.com/ under the IACUC protocol number CGUS246-CVSD24-003. Four-month-old wild type commercial cross pigs were housed in pens on raised slatted flooring in ABSL2 isolation rooms, with individual HEPA in/out air control in the animal rooms, and controlled temperature (21±2 °C) and light/dark (12 h) cycles. All pigs had *adlib* access to water and non-medicated swine finisher ration. All pigs were acclimatized for three weeks prior to start of study to ensure feed associated antimicrobial withdrawal.

### Chemical model of colitis

To induce acute colitis in mice, 6-8-week-old C57BL/6 (WT) mice were randomly exposed to dextran sodium sulfate (DSS) salt by adding 3.5% DSS (35-50 kDa, MP Biomedicals) to the drinking water for seven days. Mice were monitored daily and scored for disease activity index (DAI), which included weight change, signs of diarrhea, rectal bleeding, dehydration, and changes in behavior. Mice were assigned to control (vehicle) or experimental groups randomly; mice were treated with EcN or the GEMM BioPersist via oral gavage (0.1 mL of Luria-Bertani broth containing 1×10^9^ to 1×10^11^ CFU/mL of bacteria). Control mice were designated as the vehicle-treated group, receiving the corresponding 0.1 mL of vehicle (Luria-Bertani broth or PBS). Mice treated with 5-ASA (Sigma-Aldrich) received 75 mg/kg/day, a remission induction dose like the daily dose in humans *(27)*. Mice were anesthetized with isoflurane and sacrificed by either asphyxiation by CO_2_ or cardiac puncture, followed by cervical dislocation.

Pigs were blocked by body weight and randomly group housed in pens by sex and treatment (n= 8 male and 8 female pigs in each treatment group). To induce acute colitis pigs were administered DSS as an oral bolus of 0.125g/kg body weight in the morning on study day 0. Six hours later, each group received a version of BioPersist (63/704,043 patent pending) carrying 1×10^4^ CFU/mL of viable bacteria or an empty shell as the vehicle control. Colitis induction continued with DSS administered once daily as an oral bolus of 0.25g/kg body weight for 6 consecutive days. Clinical scores, body weight measurements, rectal temperature, and blood samples were collected on study days 0, 1, 4, and 7. Pigs were sedated with a Telazol-Ketamine-Xylazine combination (0.04mL/kg body weight; Zoetil, Virbac; Ketamine, Dechra; Rompun, Dechra) given via the intramuscular route. Thereafter pigs were euthanized with intravenous injection of pentobarbital (30mg/kg body weight; Euthanasia, VetOne). Pigs were observed for complete cessation of respiration and heartbeat, exsanguinated via jugular vein puncture and necropsy was conducted.

### Spontaneous colitis model

Five-week-old Muc2^-/-^ mice were randomly assigned to groups either control (vehicle-treated) or experimental treated with EcN or BioPersist (0.1 ml of LB containing 1×10^11^ CFU/ml of bacteria) once a week for 10-day colonization experiments or once a week for four consecutive weeks for spontaneous colitis experiments with endpoints from week 16-30. Muc2^-/-^ mice in the control group were gavaged with the vehicle (Luria-Bertani broth). Mice were monitored weekly for signs of colitis such as diarrhea, rectal bleeding, dehydration, weight change, appearance and normal behaviour. The humane endpoint included the development of rectal prolapse, requiring immediate euthanization *(34)*. Mice were anesthetized with isoflurane and sacrificed by either asphyxiation by CO_2_ or cardiac puncture, followed by cervical dislocation.

### Tissue collection

Mouse models - The small intestine, cecum, and large intestine were dissected and assessed for gross pathology. For detection of BioPerist along the GI tract, the ileum, cecum, and colon were collected separately, coiled and fixed in Carnoy’s fixative for 6 h at 4°C. For the remaining experiments, the colon was cut near the rectum, selecting the distal colon, which was cut into three sections. The most distal section was placed in 10% neutral buffered formalin and processed for histopathological analysis. The second section was stored in RNALater (Qiagen) at –80°C and designated for RNA extraction and relative qPCR. The last section was flash-frozen in liquid nitrogen and used for genomic DNA extraction. The cecum (for short-chain fatty acid analysis) and stool samples (for DNA extraction and protein markers) were immediately flash-frozen in liquid nitrogen.

Pig model – Ileum, cecum and spiral colon tissue sections and contents were collected and processed in a manner similar to the mouse samples.

### Histopathological scoring

Fixed and processed colons were embedded in paraffin and 5 µm sections were mounted in charged slides. Slides were stained for hematoxylin and eosin (Histo-core service by BCCHR, https://www.bcchr.ca/histologycore) and scored for histopathological damage by two-three observers blinded to experimental conditions using the adapted scoring systems *(34, 45, 46)*. Briefly, for the sections derived from the Muc2^-/-^ model we accounted for inflammatory cell infiltrate (0, none; 1, minimal at submucosa; 2, submucosa extending into mucosa; 3, submucosa and mucosa; 4, mucosal, submucosal and transmural), epithelial integrity (0, normal epithelium; 1, erosion; 2, crypt loss; 3, severe crypt loss; 4 severe crypt loss + ulceration), mucosal architecture (0, no irregularities; 1, irregular crypts; 2, mild hyperplasia; 3, severe hyperplasia) and edema (0, none; 1, moderate; 2, marked); for the DSS model in mice, we accounted for crypt damage (0, none; 1,basal 1/3 with mild epithelial erosion; 2, basal 2/3 with moderate epithelial erosion; 3, severe epithelial erosion with loss of goblet cells; 4, severe epithelial erosion and severe loss of goblet cells; 5, entire crypt and epithelium loss), ulceration (0, none; 1, less than 3 foci of ulceration; 2, less than 6 foci of ulceration; 3, extensive ulceration), and inflammation (0, none; 1, minimal; 2, mild; 3, moderate with mild edema; 4, marked with moderate edema; 5, severe with marked edema); for the DSS model in pigs we accounted for crypt damage (0, none; 1, hyperplastic; 2, basal 1/3 with hyperplasia; 3, basal 2/3 with moderate epithelial erosion; 4, only surface epithelium intact; 5, entire crypt and epithelium loss), ulceration (0, none; 1, less than 20 areas rich in leukocytes; 2, less than 30 areas rich in leukocytes; 3, less than 40 areas rich in leukocytes; 4, less than 50 areas rich in leukocytes and blood spots and/or abscess; 5,less than 60 areas of leukocytes and areas with extensive blood or abscesses; 6 more than 60 areas rich in leukocytes and extensive blood areas and abscesses), inflammatory cell infiltration (0, none; 1, some infiltrating cells from the submucosa into the lamina propria; 2, cells infiltrating from the submucosa into the base of the crypts plus areas of infiltration that extent further up; 3, confluent areas ∼25-50% of the mucosa infiltrated with cells; 4, confluent areas ∼50-75% of the mucosa infiltrated with cells; 5, >75% of the mucosa infiltrated with cells; 6, mucosa full of infiltrating cells), epithelial damage (0, none; 1, <10% erosion; 2, 10-25% erosion; 3, 25-50% erosion; 4, 50-75% erosion; 5, 50-75% erosion with few crypt loss; 6, extensive erosion with crypt loss; 7, complete loss of epithelium) and goblet cell depletion (0, none; 1, visible but swollen, 2, abundant goblet cells and mucus in crypts; 3, few but mucus evident in crypts; 4, some visible; 5, complete loss).

### Immunofluorescence

Colon sections were obtained from formalin-fixed tissue and embedded in paraffin. Slides were deparaffinized in xylene and then rehydrated in a gradient of ethanol. Antigen retrieval was carried out with trypsin at a concentration of 1 mg/ml at 37°C for 1 h; samples were blocked with 5% BSA-PBS for 30 min and incubated with the primary antibody (Rabbit monoclonal anti-F4/80, Cat. No. MA5-16363, and Rabbit polyclonal anti-MPO, both from Invitrogen; Rat monoclonal anti F4/80, Cat. No. CL8917 AP, Cedarlane) according to manufacturer recommendations, followed by incubation with secondary antibody (Goat anti-rat IgG AlexaFluor-conjugated 488-green antibody, Cat. No. SC-2011, Santa Cruz; Goat anti-rabbit IgG AlexaFluor-conjugated 594-red antibody, Cat. No. A11012, Invitrogen). Slides were mounted with Fluoromount containing DAPI (Invitrogen) for DNA staining of the nucleus. Images were obtained using an Olympus IX81 inverted microscope equipped with a QImaging EXi aqua Bio-Imaging camera or EVOS M5000. Images were analyzed using ImageJ. All the processing and imaging were carried out in a blind manner.

### Strain detection

#### Detection of EcN

Using primers specific to EcN (Table S3, we performed PCR on gDNA extracted from stool samples using the QIAamp DNA Stool Mini Kit or QIAamp PowerFecal DNA kit (Qiagen) with EconoTaq DNA polymerase (Lucigen) performed on the S1000 Therma Cycler (BioRad) and imaged on the UVT-28 ME gel imager Doc (Herolab).

#### Detection of *ttr* in BioPersist

gDNA was extracted from stool samples and nested PCR was performed using *ttr1* and *ttr2* set of primers (Table S3) as above. Further quantitative detection of *ttr* was performed via ddPCR with *ttr2* primers (Table S3) and QX200 EvaGreen super mix, using the QX100 droplet generator, C100 thermal cycler, QX100 droplet reader and the QuantaSoft Analysis pro software (all from BioRad). For fluorescence *in situ* hybridization (FISH) on ileum, cecum and colon, paraffin-embedded sections were deparaffinized in xylene and rehydrated in a gradient of alcohol. The sections were then incubated overnight with the probes listed in Table S3. The probes were diluted in hybridization buffer (0.9 M NaCl, 0.1 M Tris, 0.1% SDS, pH = 7.2). Then, the sections were washed once with hybridization buffer (3 min under agitation) and two more times with washing buffer (0.9 M NaCl, 0.1 M Tris, pH = 7.2). Sections were mounted in Fluormount G with DAPI and imaged with EVOS M5000 imaging system.

### qPCR analysis

RNA was extracted from colon tissue using RNeasy Fibrous Tissue Mini Kit (Qiagen). RNA derived from samples of the DSS experiment was purified using Dynabeads™ mRNA purification kit (Invitrogen). The cDNA was synthesized with iScript cDNA Synthesis Kit (BioRad). For qPCR, Sso Fast Eva Green Supermix (BioRad) and specific primers (Table S3) were used on the CFX manager 2.1 (BioRad). All the processing was carried out in a blind manner. Ct values were used to calculate the relative expression of the genes of interest, using the ΔΔCt method and normalizing gene expression to reference genes *18S, Eef2* and/or *Tbp* and relative abundance to the control group.

### Short-chain fatty acid analysis

Gas chromatography was employed to detect acetic, butyric, and propionic acid in cecal tissues as described previously *(8)*. A standard volatile acid mix (Sigma Aldrich) was used to determine the acids retention times and to construct standard curves. Data analysis was carried out in a blind manner with Chromeleon software v7.2 (Thermo Scientific), obtaining the area under the peaks and here represented as the weight percentage of the total cecal tissue (g of SCFA/g of cecal tissue x 100).

### Isolation of colonic lamina propria cells and immunophenotyping

Whole colons were collected in ice cold 1x HBSS 2% FBS. After removing the luminal content each colon was cut longitudinally and washed 3 times in fresh ice cold 1x HBSS 2% FBS and trimmed in ∼5 mm pieces; tissue was immediately transferred to a 50 ml tube containing 12 ml of EDTA-buffer (1x HBSS 2% FBS 2 mM EDTA), incubating at 37°C under agitation at 200 rpm for 15 min, repeating twice. Then tissue was rinsed three times in ice-cold RPMI-1640 and digested in 5 ml of complete media (RPMI-1640 containing 10% FBS, 1% antibiotic/antimycotic, 5mg/ml DNase I) supplemented with 5 mg/mL Liberase™ (Roche) and 5mg/ml DNase I (Sigma-Aldrich), incubating for 30 min at 37°C under agitation at 200 rpm. Digestion was stopped by adding 7 ml of ice cold EDTA-buffer. The digested tissue was passed through a 70 μM strainer and centrifuged at 400*xg* at 4°C for 10 min. The resulting cells were resuspended in complete media and proceeded to activate and/or immunophenotype. For activation, 0.1 – 0.5 x 10^6^ cells were added per well in a U-bottom 96 well plate, stimulating with leukocyte activation cocktail (BD Bioscience) and incubating for 4h at 37°C, 5% CO_2_.

Immunophenotyping for dendritic cells (DCs) and granulocytes (eosinophils, macrophages and neutrophils) only required extracellular marks, and therefore 0.5 x 10^6^ cells (previously washed in CSB buffer) were directly stained with the corresponding antibodies (listed in Table S2) in 60 μL of CSB buffer (BD Bioscience) and 10 μL of the antibody mix, incubating for 30 min at 4°C, covered from light. Stained cells were then washed (brining volume to 1 mL and centrifuging at 4°C, 4 min, 400*xg)*, viability dye 7-AAD (BD Bioscience) was added, and cells were analyzed by flow cytometry. For immunophenotyping of T cells, cells were activated, washed in PBS and stained with LIVE/DEAD dye (Thermo Fisher), then stained for extracellular marks (as above) and then fixed and permeabilized with Cyto-Fast Fix/Perm kit (BioLegend) or Mouse Foxp3 fixation and permeabilization solutions (BD Bioscience) following fabricant instructions. Cells were washed and then intracellular antibody mix was added, incubating for 20 min at room temperature and in the dark. Again, cells were washed and resuspended in CSB for flow cytometry analysis. All flow cytometry was performed with the CytoflexS (Beckman Dickinson), following the gating strategy reported in Supplementary Fig. S7) and using FlowJo v10 for final analysis. All the tissue processing and data acquisition was carried out in a blind manner.

### Bacterial translocation

Blood sample was collected from the live pig via jugular venipuncture into a sterile vacutainer with EDTA and kept on ice. Upon euthanasia, the abdominal cavity was opened, one (1) mesenteric lymph node and a small section of the spleen were collected in sterile PBS in an aseptic manner and kept on ice. Tissue samples were weighed and added to an appropriate volume of sterile PBS to make a 1:10 dilution; then macerated before further 10-fold dilutions were made. Similarly, blood samples were lysed and diluted 10-fold. All dilutions were plated on Luria-Bertani and MacConkey agar, in the absence of or with added antibiotics that were used for selection of BioPersist (Chloramphenicol 25µg/mL; Sigma Cat. No. C0378 and Kanamycin 100µg/mL; Sigma Cat. No. K1377). All agar plates were incubated at 37°C overnight and the number of colonies were counted to obtain the bacterial load for each sample type. Limit of detection was set at less than 10 colonies for the blood samples and less than 1 colony for the tissue samples.

### Statistical analyses

GraphPad Prism software v9.0 (GraphPad Software, Inc. San Diego, California, USA) was used for statistical analyses. All data was tested with D’Angostino and Pearson normality test, (alpha = 0.05). Multiple comparisons between groups were made with non-parametric Kruskal-Wallis followed by Dunn’s post-test, or One-way ANOVA followed by Tukey’s or Bonferroni’s post-test. Results are expressed as the mean ± SD, considering significant differences when *p*<0.05. Comparison of rectal prolapse rate was analyzed using a Log-rank (Mantel-Cox) test. A logistic regression model (GLM with a binomial distribution) was used to assess the probability of a positive ADG (Average Daily Gain) outcome. The binary ADG variable was dummy coded (1 for positive ADG, 0 for neutral/negative ADG). The model included “treatment” and “day” as predictors; analyses were conducted in R (v4.3.2) using the base stats package.

## List of Supplementary Materials

Fig. S1 to Fig. S8

Table S1 to Table S3

Data file S1

References *(47–52)*

## Acknowledgments

We would like to thank Dr. S. Ghosh for access to GC and A. Ames for technical assistance. We would also like to thank Dr. Jonathan Little for access to CytoflexS and Helena Neudorf for technical assistance. We would also like to acknowledge the contributions made by the co-authors AG and JAB were made while they were affiliated with UBC; co-author MKA is currently affiliated with UBC and Melius MicroBiomics.

## Funding

CONAHCYT for trainee funding (AV)

Crohn’s and Colitis Canada Innovative Grants (DLG)

Michael Smith Health Research Innovation 2 Commercialization phase 1 (DLG, LMS) Michael Smith Health Research Innovation 2 Commercialization phase 2, partnered with Melius MicroBiomics Inc. (DLG)

Melius MicroBiomics Inc. funded the porcine trial

## Author contributions

Conceptualization: AV, DLG, AG

Methodology: AV, SKG, MKA, ME, JKJ, JAB, JY, RI, NH, HM, CG

Investigation: AV, SKG, LMS, MKA, DLG

Visualization: AV

Funding acquisition: LMS, DLG

Project administration: DLG

Supervision: LMS, DLG

Writing – original draft: AV, DLG

Writing – review & editing: AV, JAB, DLG

## Data and materials availability

All data is available in this paper or in the supplemental materials. The 16S sequences files and metadata used for the analysis of this study will be uploaded to a public repository and the DOI will be made available prior to publication.

