## Supplemental figures and tables for "Bio-engineering a common probiotic to exploit colonic inflammation promotes reliable efficacy in translational models of colitis"

**BioPersist blooms under the inflammatory conditions that characterize colitis**

Muc2$^{-/-}$ start showing signs of disease as early as 8 weeks of age, but the acute phenotype characterized by diarrhea and rectal prolapse develops after 12 weeks of age. As BioPersist is a platform for a probiotic to thrive under inflammatory conditions, we decided to explore whether the onset of disease changes BioPersist load in the colon. To do this, Muc2$^{-/-}$ that received BioPersist before the onset of disease (5-8 weeks old) were monitored for the following six months (up to 30 weeks old), collecting stool samples to extract DNA and detect *ttr* copies via ddPCR. The results in Fig. S1 show that as the clinical score rises, the detected *ttr* copies increase as well, which correlates with a decrease in the clinical score for the next monitoring period. Similarly, when the clinical scores decrease, it also decreases the detection of *ttr* copies.

**BioPersist is safe**

To assess the biosafety of BioPersist in healthy mice, we administered via oral gavage 1x10$^{11}$ CFU/mL of BioPerist (one dose/day for three consecutive days), or 0.1ml of vehicle (PBS) to C57BL/6 mice. Mice were evaluated daily, recording their body weight and any signs of distress. At day 7 post-exposure to GEMMs, mice were euthanized. The colons were dissected and evaluated for macroscopic findings. The distal colon was fixed in 10% buffered formalin for 24 h and then processed, embedded and cross-sections were stained for H&E. The colon sections were evaluated for pathological damage. As shown in Fig. S2, mice receiving BioPersist have similar features as those recorded for mice receiving vehicle. The weight change (Fig. S2A) and histopathological scores and overall gross pathology (Fig. S2B) were similar among vehicle and BioPersist treated mice.

**Microbiome analysis**

To evaluate if the diversity of microbial community is altered with the intervention of BioPersist, we examined the fecal microbiome of mice exposed to BioPersist. Stool pellets were collected from Muc2$^{-/-}$ mice before and after the intervention with either vehicle, EcN or BioPersist. Pellets were flash-frozen in liquid nitrogen and stored at -80°C. Total DNA was extracted from fecal samples using QIAamp DNA stool mini kit (Qiagen) according to manufacturer's instructions. The V3-V4 region of the 16S ribosomal DNA was amplified (as described previously (*47*, *48*)) for high-throughput sequencing. Post-sequencing analyses were performed within QIIME 2 platform (*49*). qiime2R package (https://www.nature.com/articles/s41587-019-0252-6) was used for post-statistical analysis and graphic assignment in R. Paired-end sequences from two MiSeq runs were imported into QIIME 2 to perform quality-filtering, dereplication, chimera removal, denoising, and merging using the DADA2 plugin (*50*) with default settings. Data were rarefied at the depth of 19000. Exported ASV tables were merged prior to subsequent analyses. A phylogenetic tree was constructed with the q2-fragment-insertion plugin (*51*) using a backbone tree generated with the Greengene reference database (version 13_8). To minimize technical errors and noise, features with total frequency <20 or present in <3 samples were removed prior to further analysis. Taxonomic classification was collapsed and assigned at the genus level. Alpha (Shannon's diversity) and beta diversity (weighted and unweighted UniFrac) metrics were calculated with 'qiime diversity' plugin (*48*), and distance matrices were exported to R for further analysis. For Shannon diversity, we measured the changes of two time points by subtracting the initial values from the final ones. To visualize the diversity changes among the groups, ‘qiime2R’, ‘ggplot’, ‘ggpur’ packages in R v4.0.2 were used. For beta diversity, weighted and unweighted UniFrac PCoA matrices were extracted and visualized using the ‘vegan’, ‘ggplot’, ‘dplyr’ and ‘ape’ packages. The selection of two axes was determined according to the statistical summary in *Table S1* using Kruskal-Wallis test**.** A PERMANOVA ($\alpha = 0.05$) test with 999 random permutations was run on the weighted and unweighted UniFrac distance to determine the statistical significance.

After normalizing samples to their own controls, we analyzed pre- and post-treatment changes in Shannon diversity, which weighs more on rare species (Fig. S3A). We did not find significant changes in alpha diversity across groups according to the Shannon index (Fig. S3A). We aimed to look at differences in proteobacteria phylum to account for the introduction of BioPersist (Fig. S2B), and although we observed an increase in the % of proteobacteria, this change was also observed in the vehicle group. For beta diversity, we observed better separation along the timepoint with the unweighted beta diversity matrix (Fig. S3C and 3D), which primarily considers rare taxa. When analyzing weighted beta diversity plot, there was no clear group clustering (Fig. S3C). However, BioPersist group has a slightly separation between the two time points (initial vs. final) in unweighted UniFrac matrix (Fig. S3D). The results of the statistical analysis are compiled in Table S1.

**BioPersist does not protect against SHIP$^{-/-}$ ileitis**

SHIP-deficient (SHIP$^{-/-}$) mice develop spontaneous Crohn’s disease-like inflammation in the distal ileum (*22*). It starts at 4 weeks of age and is evident when mice are 6 weeks old (*22*). Pathology is driven partly by macrophage-derived IL-1β (*52*). The disease is characterized by patchy ileal thickening and redness; the histological damage includes disruption of crypt-villus architecture, immune cell infiltration, goblet cell hyperplasia and hypertrophy, ulcers, edema, and muscle thickening (*22, 52*). To assess if the GEMMs also protect against other IBD phenotypes, we introduced BioPersist in SHIP$^{-/-}$ mice. Mice heterozygous for SHIP (Inpp5d+/-) on a mixed C57BL/6x129Sv background were bred to generate SHIP$^{-/-}$ and SHIP$^{+/+}$ littermates at the Animal Research Center at BC Children’s Hospital Research Institute. When mice were 5 weeks old, they were assigned into three different groups, depending on the treatment they received: the control group received vehicle 100 uL of vehicle (LB media), BioPersist or EcN, where they received an oral gavage with $1x10^9$ CFU/ml of bacteria, for three consecutive days. When mice reached 9 weeks old, they were euthanized and the ileum (distal 15 cm), cecum and colon were collected. For the histopathological analysis, the ileum was fixed in 10% buffered formalin, processed and embdedded in paraffin, and 5 μm cross-sections were stained with H&E; the sections were scored three individuals blinded to experimental conditions. IL-1β expression was assessed in ileum homogenates via ELISA (R&D Systems, Cat. #MLB00C). For detection of BioPersist in the ileum, cecum and colon, 50-120 mg of contents were used for DNA extraction with the MagMAX Microbiome Ultra Nucleic Acid Isolation Kit (ThermoFisher Scientific, Cat #A42358) and KingFisher automated system (ThermoFisher Scientific, Cat #5400630). Neither parental (*L. reuteri*, *E. coli*) nor GEMMs (BioColonize, BioPersist) demonstrated evidence of reduced ileal inflammation characteristic in our SHIP$^{-/-}$ mouse model. SHIP$^{-/-}$ mice treated with the EcN or BioPersist showed similar gross pathology, histopathology, and histological damage scores compared to vehicle groups (Fig. S4A-C). High IL-1β concentrations are observed in full-thickness ileal homogenates of both vehicle and treated SHIP$^{-/-}$ mice (Fig. S4D). Notably, BioPersist didn’t colonize SHIP$^{-/-}$ mice. Collectively, these results suggest that the BioPersist do not reduce ileal inflammation compared to their parental strains or controls presumably because their presence is required in the gut to have an effect.

**BioPersist has a discrete effect on fecal calprotectin and secretory IgA**

Non-invasive approaches to evaluate the severity of ulcerative colitis involve the measurement of metabolites in stool samples. In human and animal models, levels of calprotectin, a protein secreted by neutrophils, correlates with inflammation and hence with disease activity for IBD. Another aspect measured in IBD patients and animal models is secretory IgA (sIgA) as it is a marker of mucosal homeostasis in the context of microbial neutralization and barrier function. In order to understand how BioPersist modulates inflammation and mucosal barrier immunity in Muc2$^{-/-}$ mice before the endpoint, either at four or six months old, we collected stool samples and measured calprotectin and sIgA via ELISA, processing the samples according to the manufacturer instructions (Abcam ab 263885 Mouse Calprotectin SimpleStep ELISA kit against S100A8/S100A9; MyBioSource MBS2702204 Mouse Secretory Immunoglobulin A (sIgA) ELISA kit). We found BioPersist treatment decreased calprotectin early (Fig. 3F), but there were no significant differences past the three months from treatment with BioPersist (Fig. S6c). For sIgA we did not find differences past the two months from treatment with BioPersist ((Fig. S6D).

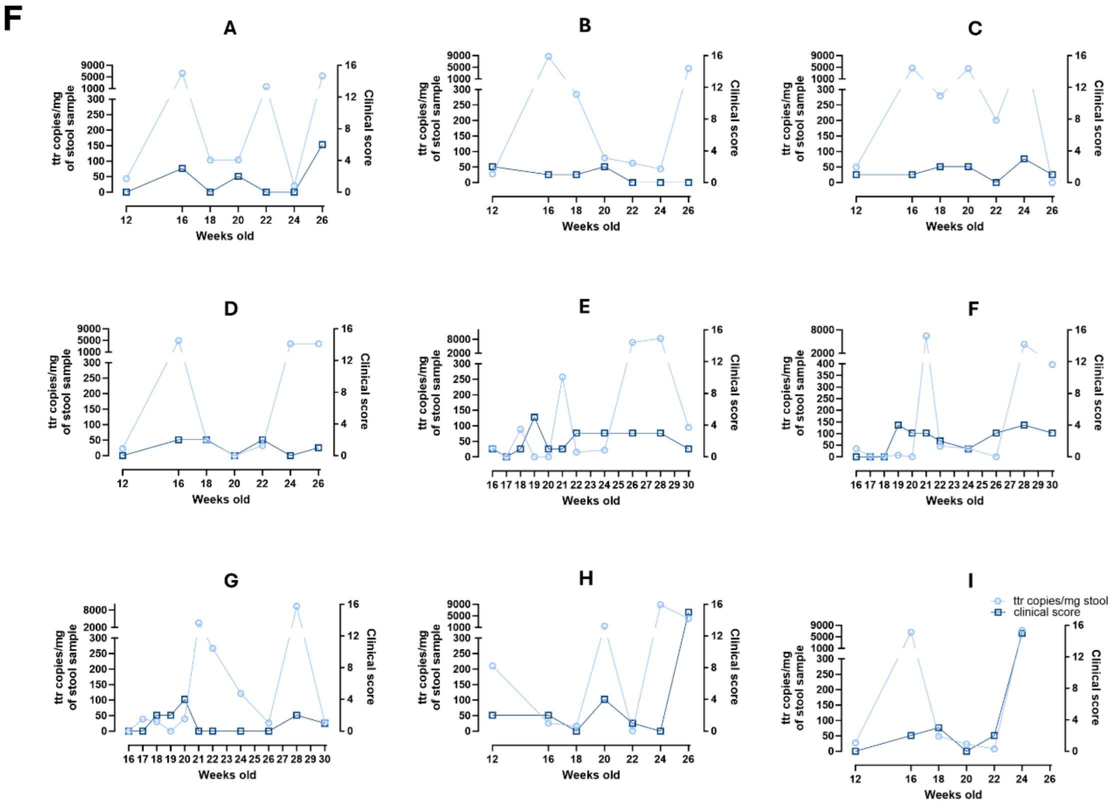


**Fig S1 Correlation between clinical scores and BioPersist detection in Muc2$^{-/-}$ mice.** A-I, Individual clinical scores for disease activity index (□, dark blue) and *ttr* loads (○, light blue) beyond 16 weeks old.

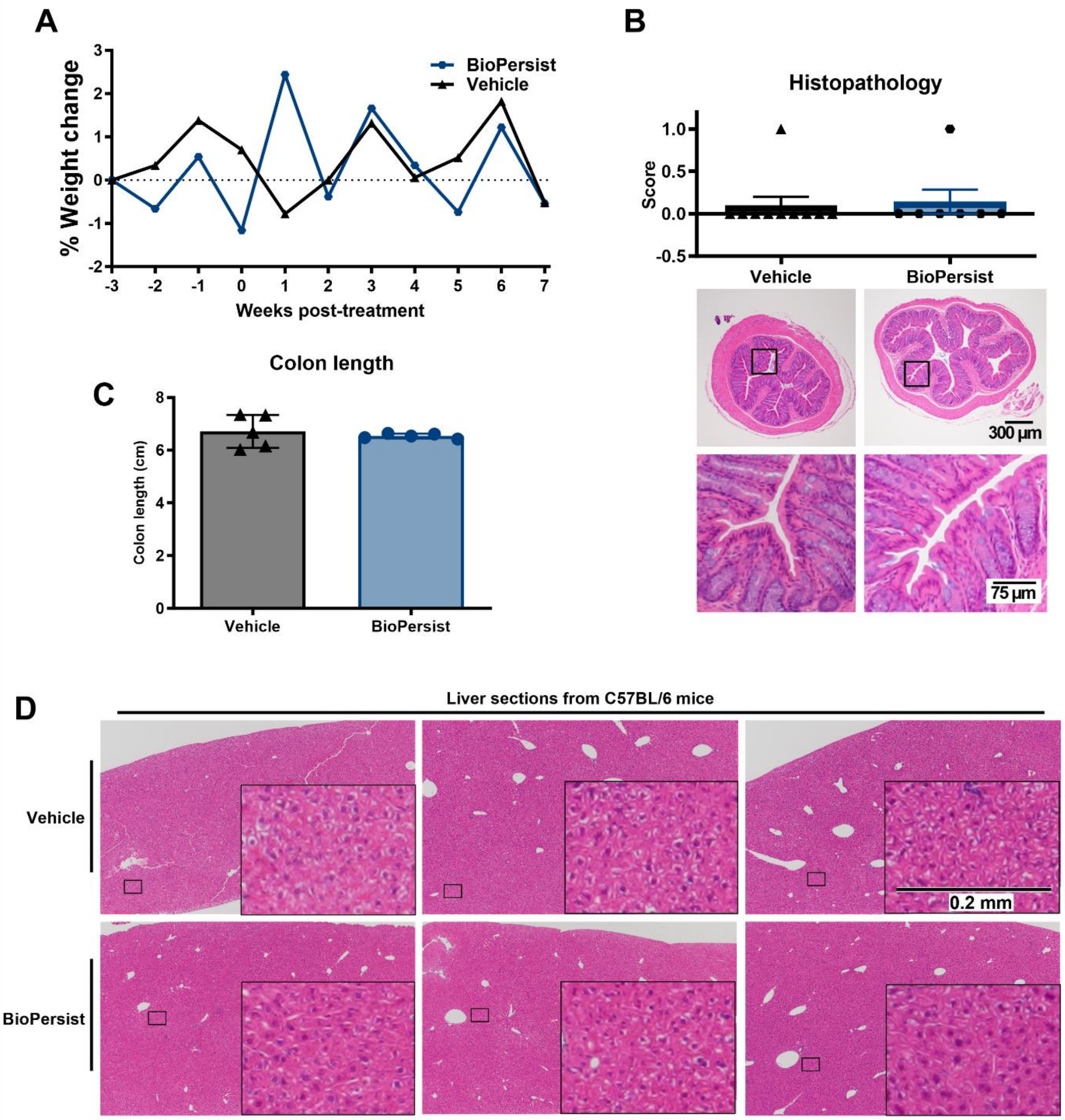


115

**Fig. S2. BioPersist in healthy mice.** A, Weight change in BioPersist and Vehicle-treated C57BL/6 mice; B, histopathological scores and representative H&E sections of the distal colon; C, colon length; D, representative liver sections of mice treated with or without BioPersist.

120

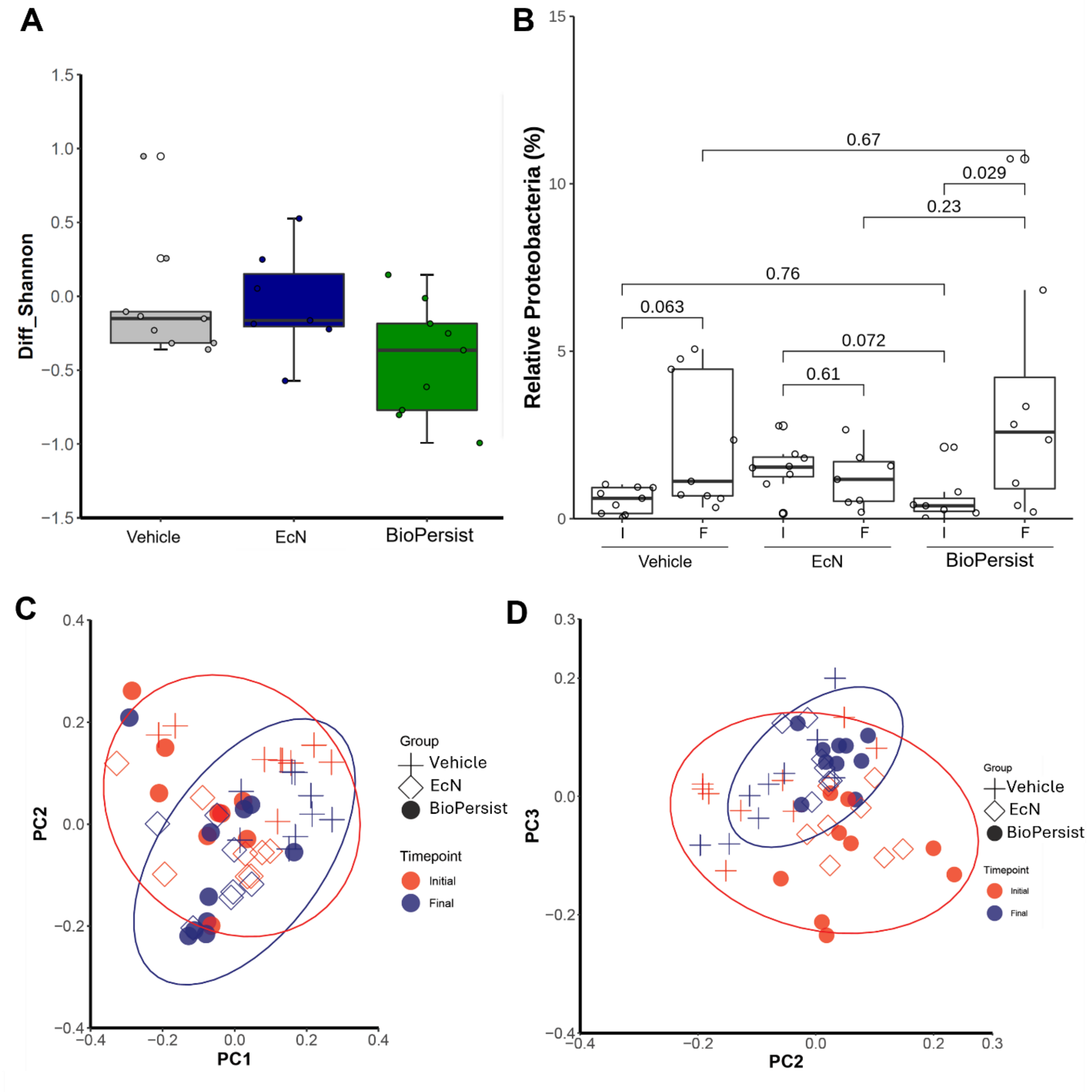


125 **Fig. S3. BioPersist does not overtake the gut microbiome**. (A) Shannon diversity shows similarities between microbial diversity across the groups. (B) % of Proteobacteria pre- (I) and post- (F) intervention across groups, along with (C) weighted and (D) unweighted beta diversity matrix shows similarities in the colonic microbiome across the groups.

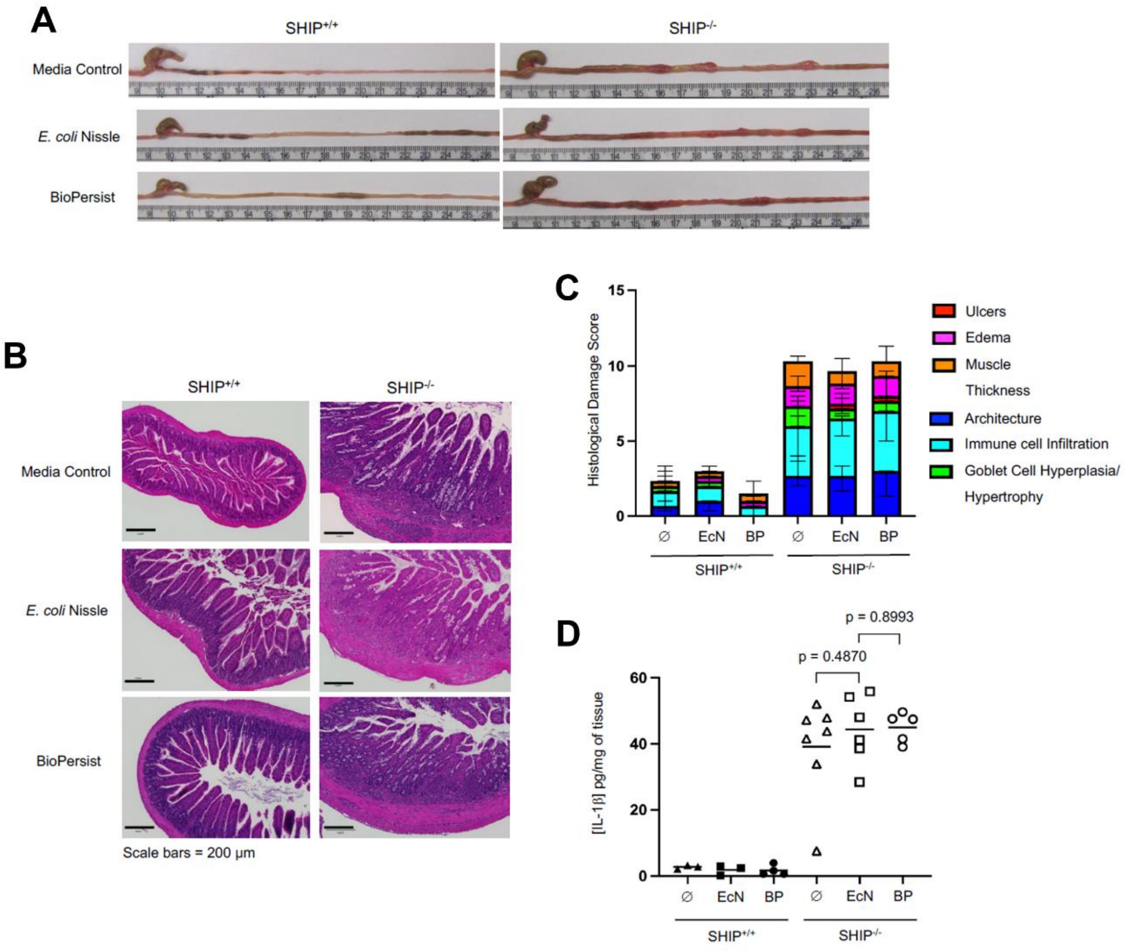


**Fig. S4. BioPersist does not reduce ileal inflammation in SHIP$^{-/-}$ mice.** A**,** Gross pathology of ilea from control and treated 9-week-old SHIP$^{+/+}$ and SHIP$^{-/-}$ mice. B**,** H&E staining of representative ileal cross-sections from untreated and treated 9-week-old SHIP$^{+/+}$ and SHIP$^{-/-}$. C, Mean histological damage scores. D, IL-1β concentrations in full-thickness ileal homogenates. No statistically significant differences were found between control (vehicle) and experimental groups using an unpaired Student's *t*-test.

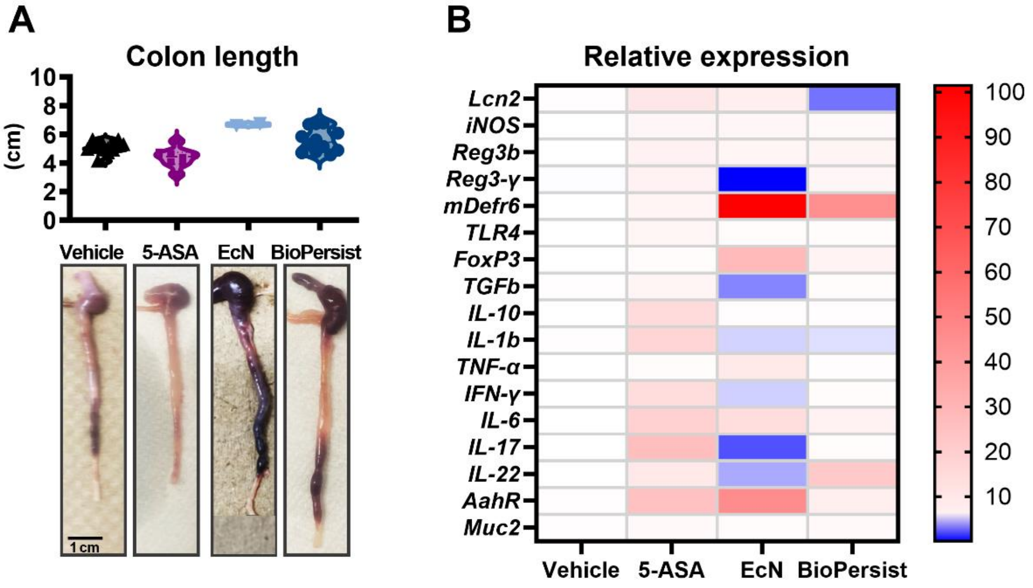


**Fig. S5. BioPersist influences visible macroscopic changes and relative changes in gene expression in mice exposed to DSS.** A**,** Gross pathology of colon derived from mice exposed to DSS. B**,** Relative gene expression in the distal colon of markers and molecules involved in barrier and/or immune function.

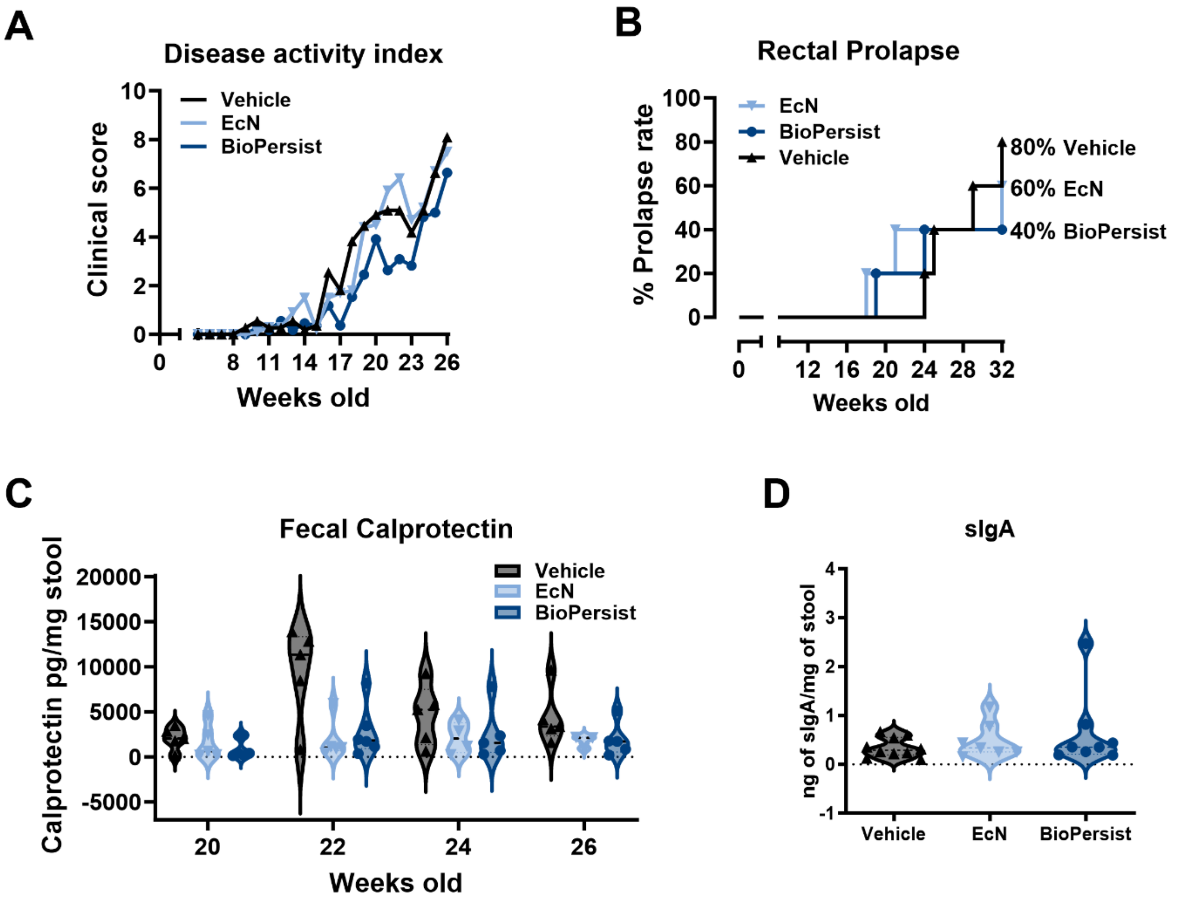


**Fig S6 BioPersist delays colitis in Muc2$^{-/-}$ mice.** A, Clinical scores of Muc2$^{-/-}$ showing the long-term endpoint beyond 16 weeks of age; B, Rectal prolapse rate; C, Fecal calprotectin quantification in stool samples; D, secretory IgA (sIgA) in stool samples of mice 16 weeks old.

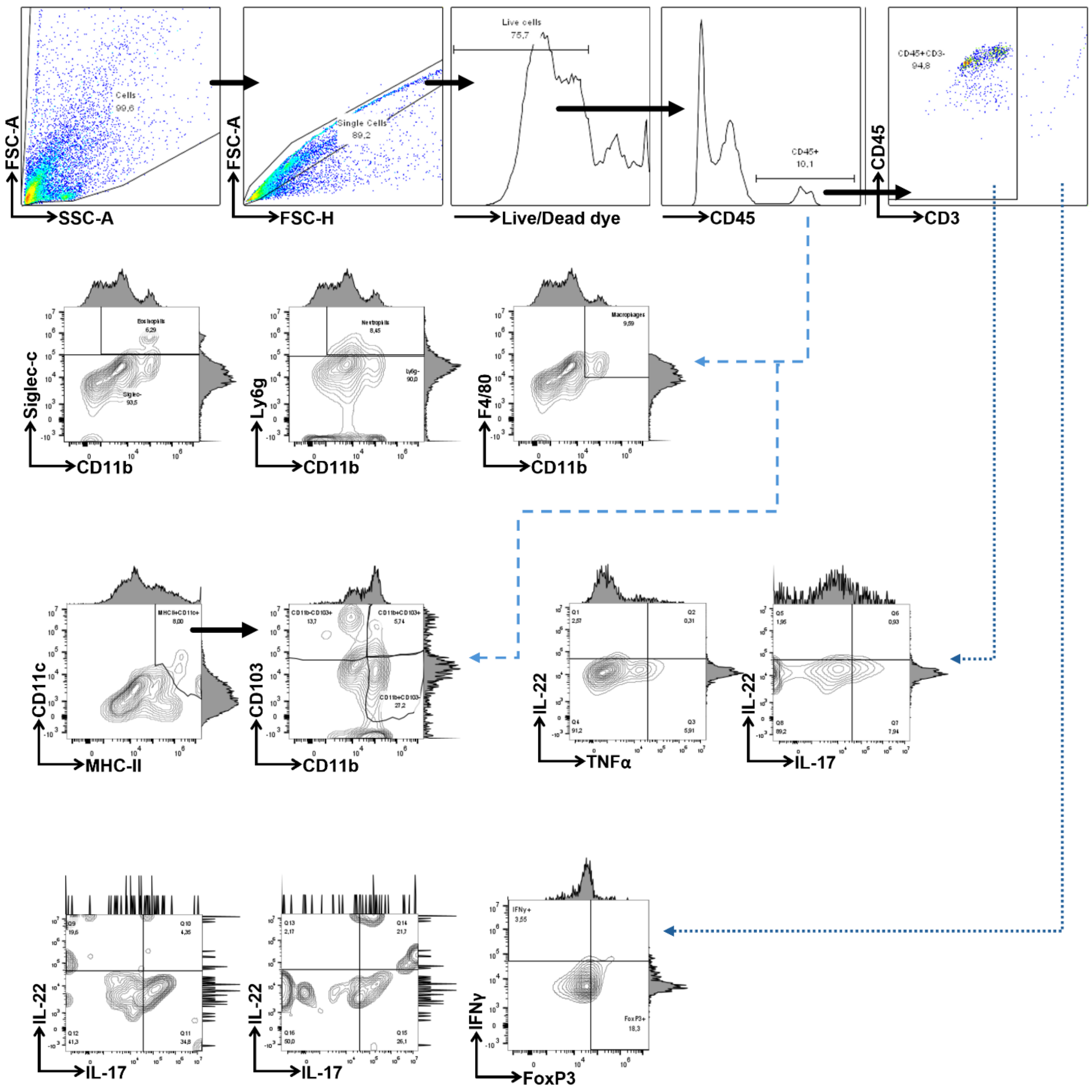


**Fig. S7. Gating strategy for lamina propria cells immunophenotyping.** Cells were sorted by size and granularity (FSC x SSC, Cells), then single cells (FSC-H x FSC-A, Singlet cells), live/dead exclusion (Live/Dead dye, Live cells), and those positive for leukocyte marker CD45 (CD45+). After this, cells were sorted for granulocyte markers (CD11b, Ly6g, Siglec-F, and F4/80), dendritic cell markers (MHC-ii, CD11c, CD11b, and CD103), and lymphocyte markers (CD3, FoxP3) and selectec cytokines (IFNγ, IL17, IL-22, TNFα). Data was collected on a CytoflexS and the results were analyzed with FlowJo v10.10.

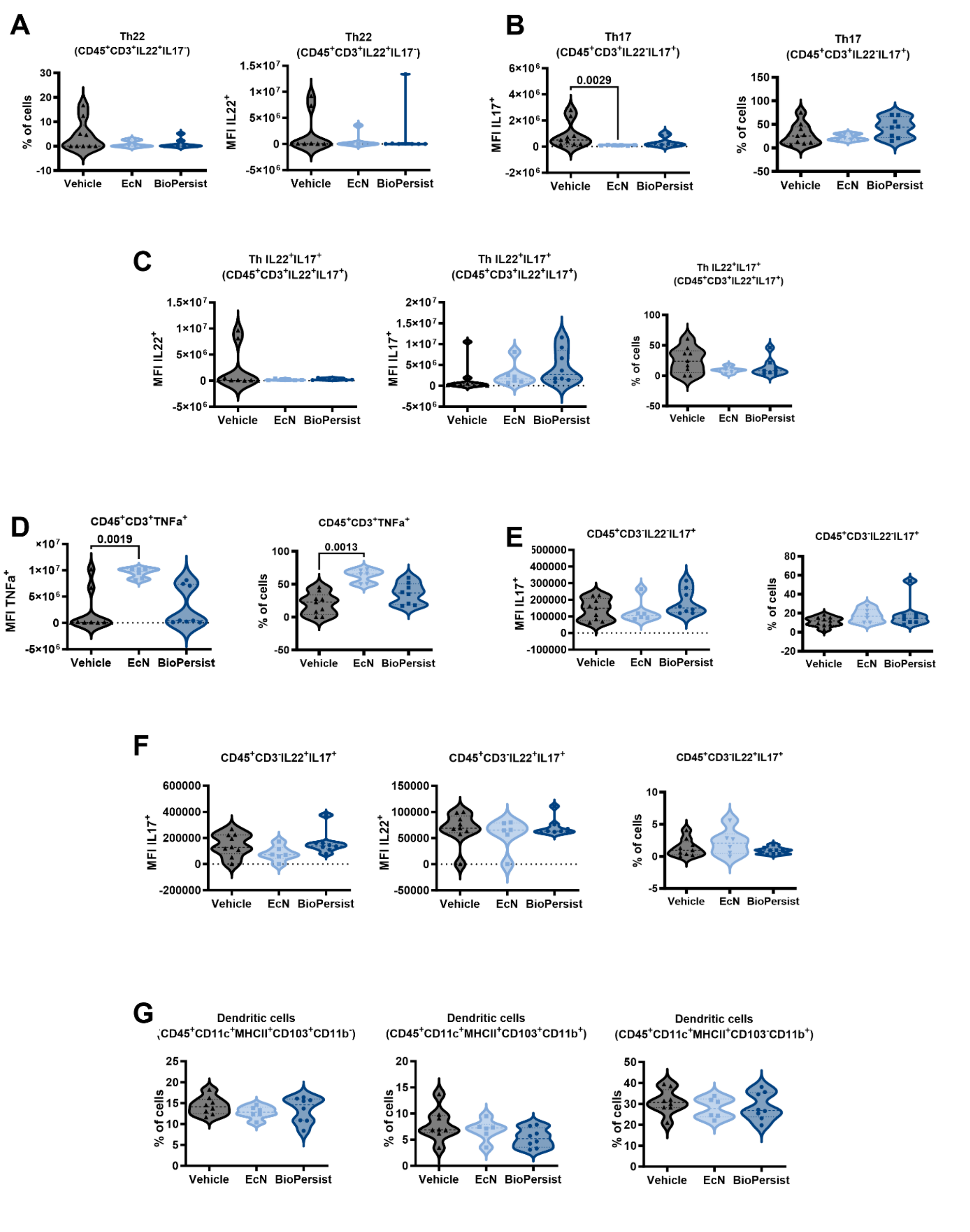


**Fig. S8. Immune cells not affected by BioPersist.** A – G show different immune cells either T cells (CD3+, panels A-D) or leukocytes that are CD3- (E and F), as well as dendritic cells (G). Data was analyzed with Kruskal-Wallis (A-G). MFI, median fluorescence intensity.

**Table S1. Statistical summary for microbiome analysis**

| | Group1 | Group2 | p-value | p.adj | p.signif | Method |
|---|---|---|---|---|---|---|
| **Shannon** | Vehicle | EcN | 0.673 | 0.67 | ns | Wilcoxon |
| | Vehicle | BioPersist | 0.094 | 0.19 | ns | |
| | EcN | BioPersist | 0.046 | 0.14 | ns | |

| | Axis | p-value | p.adj | p.signif | Method |
|---|---|---|---|---|---|
| **Weighted_Unifrac** | PC1 | 0.0015 | 0.0015 | ** | Kruskal-Wallis |
| | PC2 | 0.0005 | 0.00045 | *** | |
| | PC3 | 0.0179 | 0.018 | * | |
| | PC4 | 0.0227 | 0.023 | * | |
| | PC5 | 0.0589 | 0.059 | ns | |
| | PC6 | 0.369 | 0.37 | ns | |
| **Unweighted_Unifrac** | PC1 | 0.0805 | 0.08 | ns | Kruskal-Wallis |
| | PC2 | 0.0019 | 0.0019 | ** | |
| | PC3 | 0.0002 | 0.00023 | *** | |
| | PC4 | 0.05 | 0.05 | ns | |
| | PC5 | 0.0379 | 0.038 | * | |
| | PC6 | 0.591 | 0.59 | ns | |

| | | | **PC1** | | **PC2** | |
|---|---|---|---|---|---|---|
| | **Group1** | **Group2** | p-value | p.signif | p-value | p.signif |
| **Weighted Unifrac Multi-comparison Summary** | Vehicle-I | Vehicle-F | 0.258 | ns | 0.001 | ** |
| | EcN-I | EcN -F | 0.463 | ns | 0.281 | ns |
| | BioPersist-I | BioPersist -F | 0.968 | ns | 0.095 | ns |
| | Vehicle -I | EcN -I | 0.036 | * | 0 | *** |
| | Vehicle-I | BioPersist -I | 0.019 | * | 0.063 | ns |
| | EcN-I | BioPersist -I | 0.236 | ns | 0.114 | ns |
| | Vehicle -F | EcN -F | 0.001 | *** | 0.008 | ** |
| | Vehicle -F | BioPersist -F | 0.001 | ** | 0.065 | ns |
| | EcN-F | BioPersist-F | 0.669 | ns | 0.887 | ns |

| | | | **PC2** | | **PC3** | |
|---|---|---|---|---|---|---|
| | **Group1** | **Group2** | p-value | p.signif | p-value | p.signif |
| **Unweighted Unifrac Multi-comparison Summary** | Vehicle-I | Vehicle-F | 0.667 | ns | 0.931 | ns |
| | EcN-I | EcN -F | 0.054 | ns | **0.004** | ** |
| | BioPersist-I | BioPersist -F | 0.72 | ns | **3.00E-04** | *** |
| | Vehicle -I | EcN -I | 0.011 | * | 0.139 | ns |
| | Vehicle-I | BioPersist -I | 0.019 | * | 0.006 | ** |
| | EcN-I | BioPersist -I | 0.888 | ns | 0.2 | ns |

| | | | | | |
|---|---|---|---|---|---|
| Vehicle - F | EcN -F | 0.174 | ns | 0.252 | ns |
| Vehicle - F | BioPersist -F | 0.003 | ** | 0.133 | ns |
| EcN-F | BioPersist-F | 0.088 | ns | 0.813 | ns |

**Table S2. Antibodies used**

| Antibody | Fluorophore | Clone | Vendor | Catalog Number |
|---|---|---|---|---|
| Rat anti-mouse CD103 | PE | M290 | BD Biosciences | 557495 |
| Rat anti-mouse CD11b | FITC | M1/70 | BD Biosciences | 553310 |
| Hamster anti mouse CD11c | APC | HL3 | BD Biosciences | 550261 |
| Rat anti-mouse CD3 | PerCP Cy5.5 | 17A2 | BD Biosciences | 560527 |
| Rat anti-mouse CD4 | Pacific Blue | RM4-5 | BD Biosciences | 558107 |
| Rat anti-mouse CD45 | PE/Cy7 | 30-F11 | Biolegend | 103114 |
| Rat anti- mouse F4/80 | PerCP Cy5.5 | BM8 | Biolegend | 123128 |
| Rat anti-mouse Foxp3 | PE | R16-715 | BD Biosciences | 563101 |
| Rat anti-mouse IFNy | PerCP Cy5.5 | XMG1.2 | BD Biosciences | 560660 |
| Rat anti-mouse IL-17A | PE | TC11-18H10 | BD Biosciences | 559502 |
| Rat anti-mouse Ly6G | PE | 1A8 | Biolegend | 551461 |
| Rat anti-mouse MHC-II | Pacific Blue | M5/114.15.2 | Biolegend | 107620 |
| Rat Anti-Mouse Siglec-F | BV421 | E50-2440 | BD Biosciences | 562681 |
| Rat anti-mouse TNF | APC | MP6-XT22 | BD Biosciences | 554420 |
| Rabbit monoclonal anti F4/80 | | | Invitrogen | MA5-16363 |
| Rabbit polyclonal anti-MPO | | | Invitrogen | PA5-16672 |
| Rat monoclonal anti F4/80 | | | Cedarlane | CL8917 AP |
| Goat anti-rat IgG | AF488 | | Santa Cruz | Sc-2011 |
| Goat anti-rabbit IgG | AF594 | | Invitrogen | A11012 |

**Table S3. Primers and probes used**

| **Target** | **Sequence (5' → 3')** | **Tm** |
|---|---|---|
| EcN1F | TGACGATGCCTTAGATCTTGATGC | 59 |
| EcN1R | GCTTCTTGTATATGGTTATACAATTGTCGTC | 59 |
| EcN2F | GACAGTGAGCTAAAACAGTCC | 56 |
| EcN2R | GAAATGGTGTTGGTGCGGTC | 60 |
| ttr1 F | CAGAGCGAGAAATCCGGTAA | 55 |
| ttr1 R | GGTAGTCAGGAAGTCACGAATG | 57 |
| ttr2 F | ATTCAGACCTCCTGCCAAAG | 57 |
| ttr2 R | CCTTACGACGCCCGATTTAT | 58 |
| ttr2 -TxRed | ATTCAGACCTCCTGCCAAAG | 57 |
| ttrS - TxRed | GCAACCGTTGGCAAAGACAT | 60 |
| ttr2_qR | ACGTCGCCCCATGTTTTCTA | 56 |
| 18s F | CGGCTACCACCCAAGGAA | 58 |
| 18S R | GCTGGAATTACCGCGGCT | 58 |
| EEF2 F | TGTCAGTCATCGCCCATGTG | 58 |

| | | |
|---|---|---|
| EEF2 R | CATCCTTGCGAGTGTCAGTGA | 58 |
| Tbp F | ACCGTGAATCTTGGCTGTAAAC | 58 |
| Tbp R | ACCGTGAATCTTGGCTGTAAAC | 58 |
| Muc2 F | GCCAGATCCCGAAACCA | 58 |
| Muc2 R | TATAGGAGTCTCGGCAGTCA | 58 |
| TNFalpha F | CATCTTCTCAAAATTCGAGTGACAA | 58 |
| TNFalpha R | TGGGAGTAGAACAAGGTACAACCC | 58 |
| IFNy F | TCAAGTGGCATAGATGTGGAAGA | 58 |
| IFNy R | TGGCTCTGCAGGATTTTCATG | 58 |
| IL-17 F | TCCCTCTGTGATCTGGGAAG | 58 |
| IL-17 R | CTCGACCCTGAAAGTGAAGG | 58 |
| IL-10 F | AGGGCCCTTTGCTATGGTGT | 58 |
| IL-10 R | TGGCCACAGTTTTCAGGGAT | 58 |
| Reg3y F | CCCGTATAACCATCACCATCAT | 58 |
| Reg3y R | GGCATCTTTCTTGGCAACTTC | 58 |
| RELMbeta F | ATGGGTGTCACTGGATGTGCTT | 58 |
| RELMbeta R | TGGGAGTAGAACAAGGTACAACCC | 58 |
| IL-22 F | AGCTCCTGTCACATCAGCG | 58 |
| IL-22 R | AGCTTCTTCTCGCTCAGACG | 58 |
